# *SerraNA*: a program to determine nucleic acids elasticity from simulation data

**DOI:** 10.1101/2020.03.24.004945

**Authors:** Victor Velasco-Berrelleza, Matthew Burman, Jack W. Shepherd, Mark C. Leake, Ramin Golestanian, Agnes Noy

## Abstract

The resistance of DNA to stretch, twist and bend is broadly well estimated by experiments and is important for gene regulation and chromosome packing. However, their sequence-dependence and how bulk elastic constants emerge from local fluctuations is less understood. Here, we present *SerraNA*, which is an open software that calculates elastic parameters of double-stranded nucleic acids from dinucleotide length up to the whole molecule using ensembles from numerical simulations. The program reveals that global bendability emerge from local periodic bending angles in phase with the DNA helicoidal shape. We also apply *SerraNA* to the whole set of 136 tetra-bp combinations and we observe a high degree of sequence-dependence for all elastic parameters with differences over 200%. Tetramers with TA and CA base-pair steps are especially flexible, while tetramers containing AA and AT tend to be the most rigid. Our results thus suggest AT-rich motifs generate extreme mechanical properties depending of the exact sequence ordering, which seems critical for creating strong global bendability on longer sequences when phased properly. *SerraNA* is a tool to be applied in the next generation of interdisciplinary investigations to further understand what determines the elasticity of DNA.

**Graphical TOC Entry:** 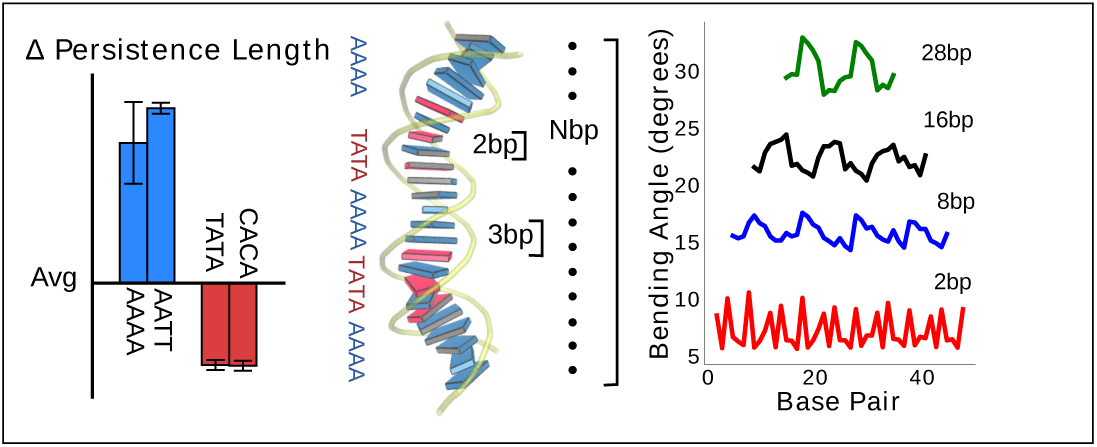

## Introduction

For genomes to function properly, chromosomes need to fold into a hierarchy of structures, causing, for example, expression correlation of genes located within the same topological domain.^1^ Besides, it is widely known that DNA looping is a fundamental structure for gene regulation that facilitates long-range communication between a promoter and its distal regulatory elements.^2,3^ Moreover, DNA can be subjected to forces up to tens of pN approximately in cells due to the activity of protein motors.^4^ And finally, on the shortest scale, DNA distortion has been detected as determining the formation of diverse DNA:protein complexes like nucleosomes, some transcription factors or bacterial nucleoid association proteins.^5^ Therefore, it is important to measure the mechanical response of DNA to bending, stretching and torsion, which is well established to have average values close to 50 nm for the persistence length,^6–10^ between 1100-1500 pN for the stretch modulus^11,12^ and ranging from 90 to 120 nm for the torsion elastic constants^8,13,14^ (for a good summary of experimental values see Lipfert *et al.*^15^).

What is less clear from experimental data is the spread of elastic properties depending on sequence and which local elements build up the bulk flexibility of long DNA fragments. There have been several attempts to deduce the particular values associated to a sequence from cyclization probabilities, although these methodologies are not unambiguous and require the use of theoretical models.^16,17^ In addition, it has been very difficult to identify the mechanisms through which some short sequence motifs, like A-tracts, originate extraordinary bending.^18,19^ On these matters, molecular dynamics (MD) simulations at atomic resolution have become an impressive source of new important information,^20^ that have provided (i) systematic analysis at the dinucleotide level,^21,22^ an evaluation of the influence of nearest flanking base-pairs (bp) up to the tetranucleotide level^23,24^ and, among others, (iii) an explanation of contradictory stiffness data on A-tracts.^25^ On a more coarse-grained level, Monte Carlo (MC) simulations have found that most of sequence-dependence variability is originated at the level of static curvature.^26^

Previously, we designed the Length-Dependent Elastic Model (LDEM) for describing how bulk elastic properties emerge from bp fluctuations using the sampling obtained by nucleic acids simulations.^27^ The LDEM revealed that the crossover from local to global occurs typically within one helical turn of DNA^27^ as has been confirmed by others.^28,29^ In terms of torsion elasticity, we observed a transition from dinucleotide values of 30-50 nm to the long-range elastic constants of 90-120 nm in agreement with experimental data.^8,13,14,30^ The model also revealed that stretch modulus changed as a function of molecular length in a non-monotonic way on shorter scales followed by a stabilization to similar values of force-extension measurements (1100-1500 pN).^11,12,27,29^ Highly soft stretch modulus measured by SAXS experiments on short oligomers31 was observed to be caused mainly by end effects.^27^ For the persistence length, we found that the periodic tangent-tangent correlation reflected the “crookedness”^32^ of the static curvature of the DNA helix^26,27,29,30,32–35^and, without considering these modulations, the decay was close to the consensus value of 50 nm.^6–10^ Thus, the LDEM is suitable for describing the average mechanical properties of DNA and, from this perspective, it was applied to test the DNA force-field for atomic simulations, Parmbsc1.^36^

Here we present *SerraNA*, which is an open-source, versatile and integrated implementation of the LDEM, that allows fast simulation analysis and detection of emergent sequence effects. It calculates the overall elastic constants of helical nucleic acids (NA) and the elastic/structure profiles for every possible sub-length (*serra* from Latin means “mountain range”). To our knowledge, there is no other program that estimates bulk flexibility constants from ensembles obtained by numerical simulations and that uncovers systematically how these properties emerge from local sequence-dependence fluctuations.

The paper describes the theoretical background behind the LDEM and it provides estimations for the different elastic constants by using MD simulations over a series of DNA fragments between 32 to 62 bp. Then, *SerraNA* is used to determine how bulk elastic constants emerge from local fluctuations using bendability as an example. Finally, the program is applied to the ABC trajectory database,^23^ which contains all the possible tetra-bp combinations, for exhaustively evaluating the dependence on sequence of the elasticity of DNA.

## The Length-Dependent Elastic Model (LDEM)

### Geometric description for different fragments lengths

The two bending angles, roll and tilt, and the rotational angle twist at the bp-step level are adapted to evaluate the relative orientation of a pair of bp spaced by an increasing number of nucleotides. The vertical displacement, which is associated with stretch, is characterized by end-to-end distance but a fragment’s contour length is also calculated for a more comprehensive description of the polymer structure (see below and Figure 1 for further details).

**Figure 1:**
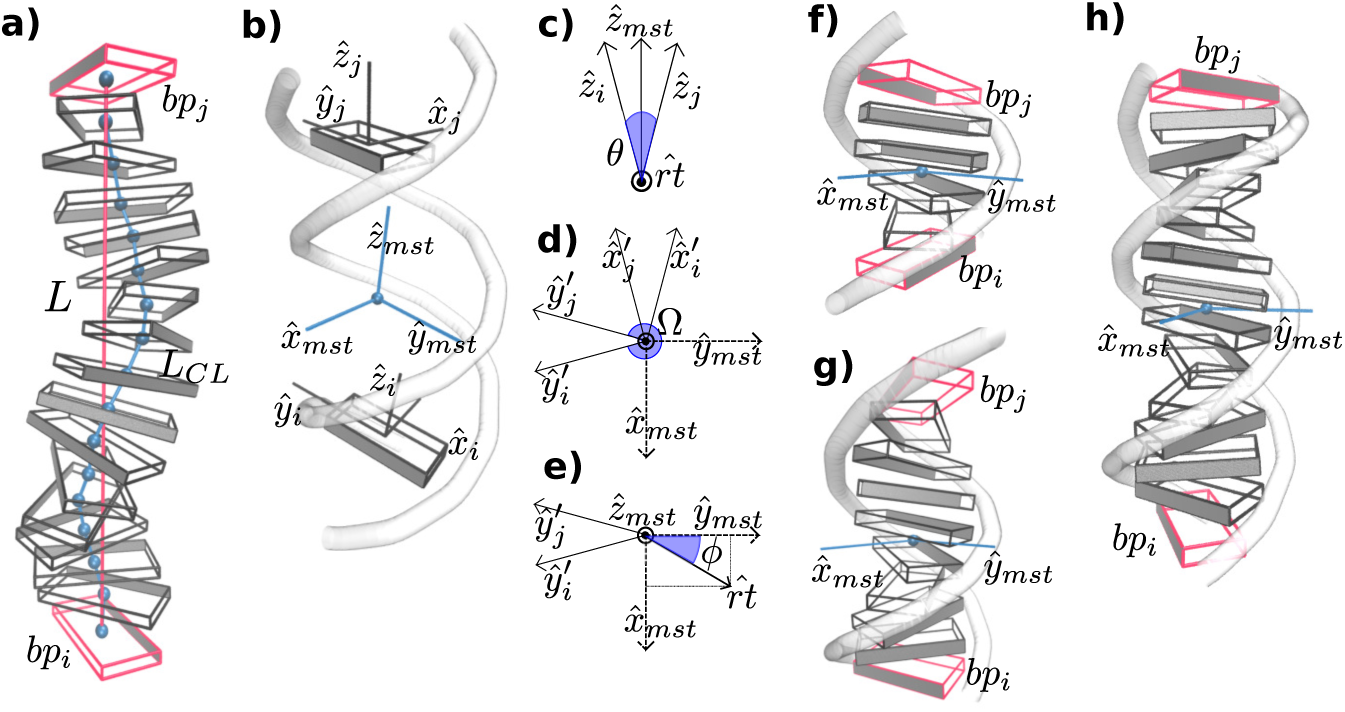
Schematic diagrams of the algorithm implemented in *SerraNA* for calculating the geometric parameters at different fragment lengths. (a) Vertical displacement is characterize by end-to-end distance (in red) and contour length (in blue). (b) Twist and bend angles between bp *i* and *j* are obtained via the mid-base triad (T_*mst*_) positioned at the mid-point. (c) Bending angle *θ* and bending axis 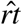 are defined by directional vectors 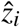 and 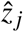. Co-planar vectors *ŷ*′_*i*_, *ŷ*′_*j*_, 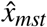 and *ŷ*_*mst*_ define twist angle Ω (d) and roll and tilt bending angles (e). (f)-(h) T_*mst*_ between bp (in red) separated by 4, 8 and 12 bp, respectively. Roll and tilt consist on bending towards grooves and backbone, respectively, at the fragment midpoint for the different sub-fragment lengths. Angles and T_*mst*_ are highlighted in blue.

The spatial configuration of a bp *i* is specified by giving the location of a reference point 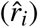 and the orientation of a right-handed orthonormal reference triad (T_*i*_) following the mathematical procedure of the 3DNA program,^37^ where *ŷ*_*i*_ points to the backbone of first strand, 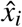 points to the major groove and 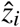 marks the molecular direction at that particular point (Figure 1b). Then, the CEHS scheme is applied for obtaining the molecular twist and the roll/tilt contributions to bend38,39 (Figure 1). The algorithm is used to calculate the mid-step triad T_*mst*_ between bp *i* and *j* that define an oligomer whose length ranges from 2 bp to N (Figure 1b). N is the total number of bp in the DNA fragment minus the two for each end, which have been discarded in order to avoid temporary loss of base pairing and other end effects.

The bending angle *θ* is obtained directly from the direction correlation 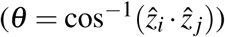 and the corresponding bending axis 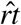 is calculated by 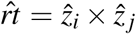 (Figure 1b). Next, T_*i*_ and T_*j*_ are rotated around 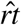 by half of *θ* for obtaining T′_*i*_ = R_*rt*_ (+*θ*/2)T_*i*_ and T′_*j*_ = R_*rt*_ (−*θ*/2)T_*j*_, where the transformed *x*-*y* planes are now parallel with each other and their *z*-axes coincide (see Figure 1c and 1d). T_*mst*_ is directly built by averaging and normalizing T′_*i*_ and T′_*j*_. The corresponding 3 rotations (tilt *τ*, roll *ρ*, twist Ω) are defined as:

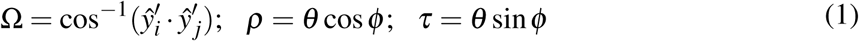

where *φ* is the angle between 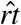 and the *ŷ*_*mst*_ (Figure 1e). Note that roll and tilt variables in lengths longer than a dinucleotide denote bending towards grooves and backbone direction, respectively, according the T_*mst*_ i.e. the fragment midpoint (see Figure 1).

For each DNA sub-fragment, end-to-end distance (*L*) and contour length (*L*_*CL*_) are defined as:

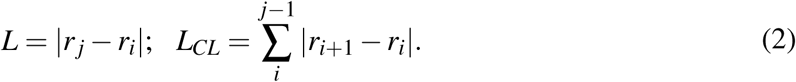

For completeness, the three rigid-body translation variables at the dinucleotide level (shift *X*_*i,i*+1_, slide *Y*_*i,i*+1_ and rise *Z*_*i,i*+1_) are calculated by:

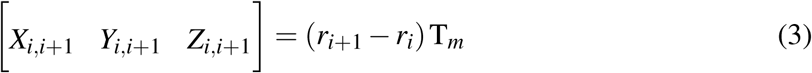

and the extrapolation to longer scales can be designated by:

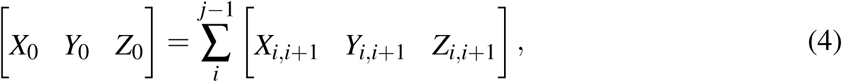

where added-shift *X*_0_, added-slide *Y*_0_ and added-rise *Z*_0_ can be interpreted structurally as the three pseudo components of *L*_*CL*_.

For better comparison with experiments, only end-to-end distance *L*, twist Ω, roll *ρ* and tilt *τ* are utilized for the calculation of DNA elastic constants. *SerraNA* outputs the ‘structural_parameters.out’ file with the complete set of structural variables (including total bending angle, directional correlation, contour length and added shift, slide and rise) at all lengths with the idea of providing a full conformational illustration of the whole molecular stretch (see flowchart in Figure 2 for more details).

**Figure 2:**
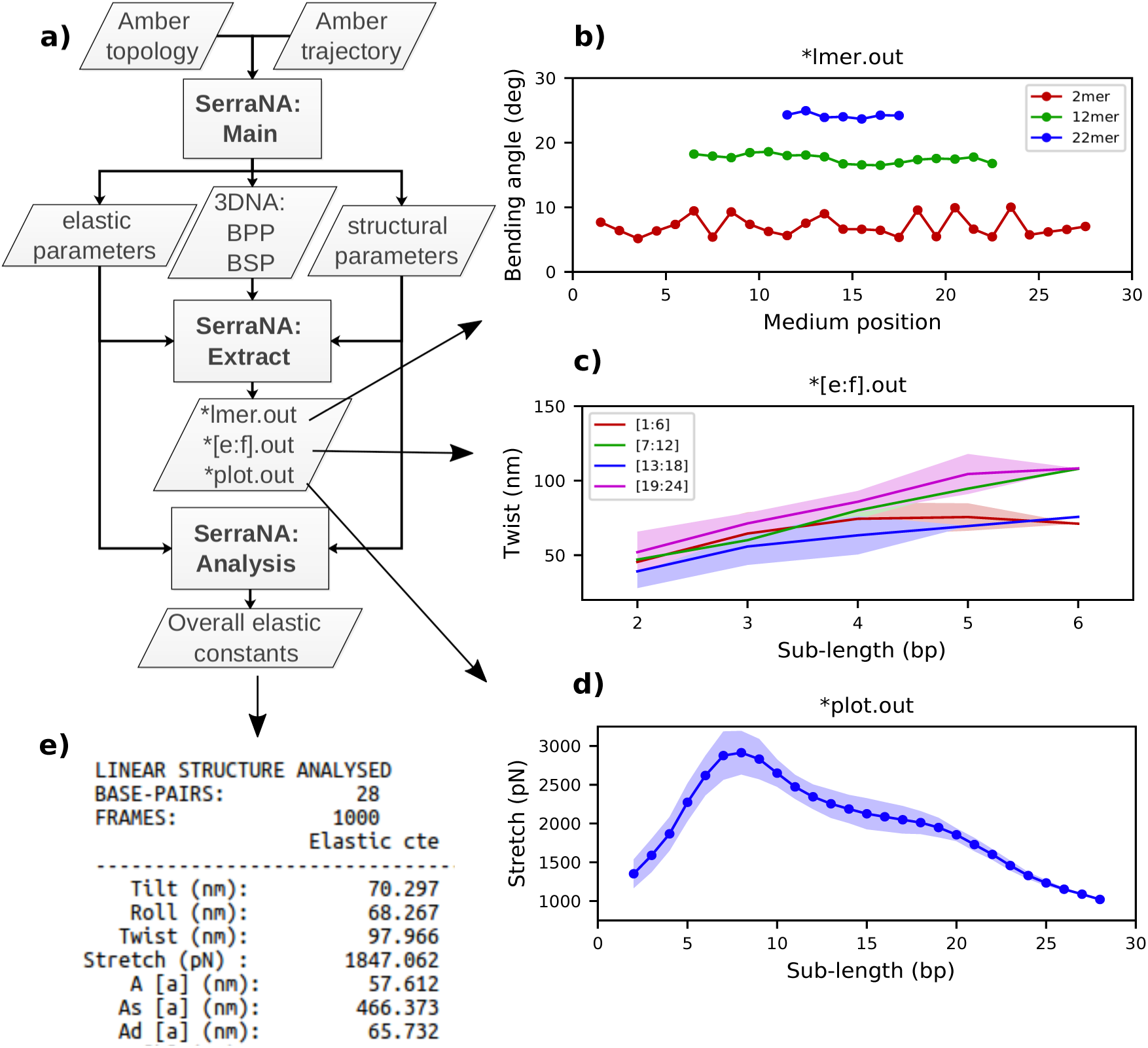
General workflow of *SerraNA* using 32mer as an example. (a) Main program outputs bp and bp-step parameters (BPP and BSP, respectively), together with structural and elastic parameters at different lengths. *Extract* tool creates simple files for (b) plotting profiles along the molecule for a sub-length l (*lmer.out) and for (c) plotting the length-dependence from bp e to f (*[e:f].out) or (d) from the whole fragment (*plot.out). (e) *Analysis* tool extracts the overall elastic constants from a NA molecule.

### The length-dependent model of DNA elasticity

Under the assumption that distribution of values adopted by a variable *X* is fully Gaussian and non-correlated with the rest of deformation parameters, the corresponding elastic constant *K* for a particular length can be easily derived from its variance *Var*(*X*) estimated during a MD trajectory:^27,40^

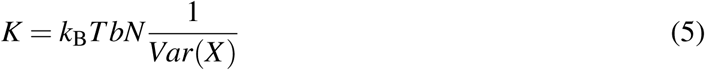

where fragment length or sub-lengths are specified by *N* dinucleotide steps with rise *b* = 0.34 nm. However, the four distortion variables chosen to describe DNA flexibility on this model (roll, tilt, twist and end-to-end) are non-orthogonal. This effect is specifically taken into consideration by determining elastic constants as the diagonal terms of the inverse covariance matrix *V*^−1^ or elastic matrix *F*:^41^

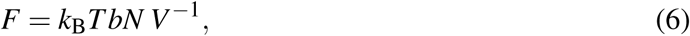

Correspondingly, the diagonal terms of *V*^−1^ can be understood as the reciprocal of the partial variances, (1/*Var*_*p*_(*X*)). *Var*_*p*_(*X*) is a measure of the residual variance associated with a deformation after removing the linear effects caused by other variables.^27,42^ All terms from the different *F*s calculated using all possible sub-fragments are printed in the ‘elastic_parameters.out’ output file for a complete dynamic description of the NA molecule (see Figure 2).

### Estimation of bulk twist elastic constant

The twist elastic constant for a singular sub-fragment *k* (*C*_*k*_) is the diagonal term of *F*_*k*_ corresponding to twist, *F*_*k*_ being the elastic matrix associated to that particular DNA sub-fragment. Then, the twist elastic modulus as a function of length (*C*_*N*_) is calculated by averaging all sub-fragments *k* with the same number of dinucleotide steps *N*: *C*_*N*_ = 〈*C*_*k,N*_〉 (Figure 3). Because the transition from bp level to the global elastic behavior occurs within one helix turn, values at lengths longer than 12 bp can already be considered good estimations of bulk twist elastic modulus *C* (Figure 3a). Global *C* of an individual DNA fragment is calculated as the overall average of the series of *C*_*N*_:

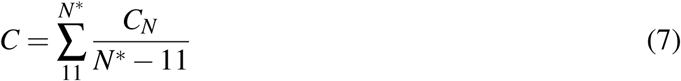

where *N* ranges from 11 bp-steps to *N∗*, *N∗* being the maximum sub-fragment length considered. By default *SerraNA* discards the ten longest sub-fragment lengths for counting *N∗* in order to have at least ten different values in averaging *C*_*N*_, but this is an option that can be modified in the program (see Figure 2).

**Figure 3:**
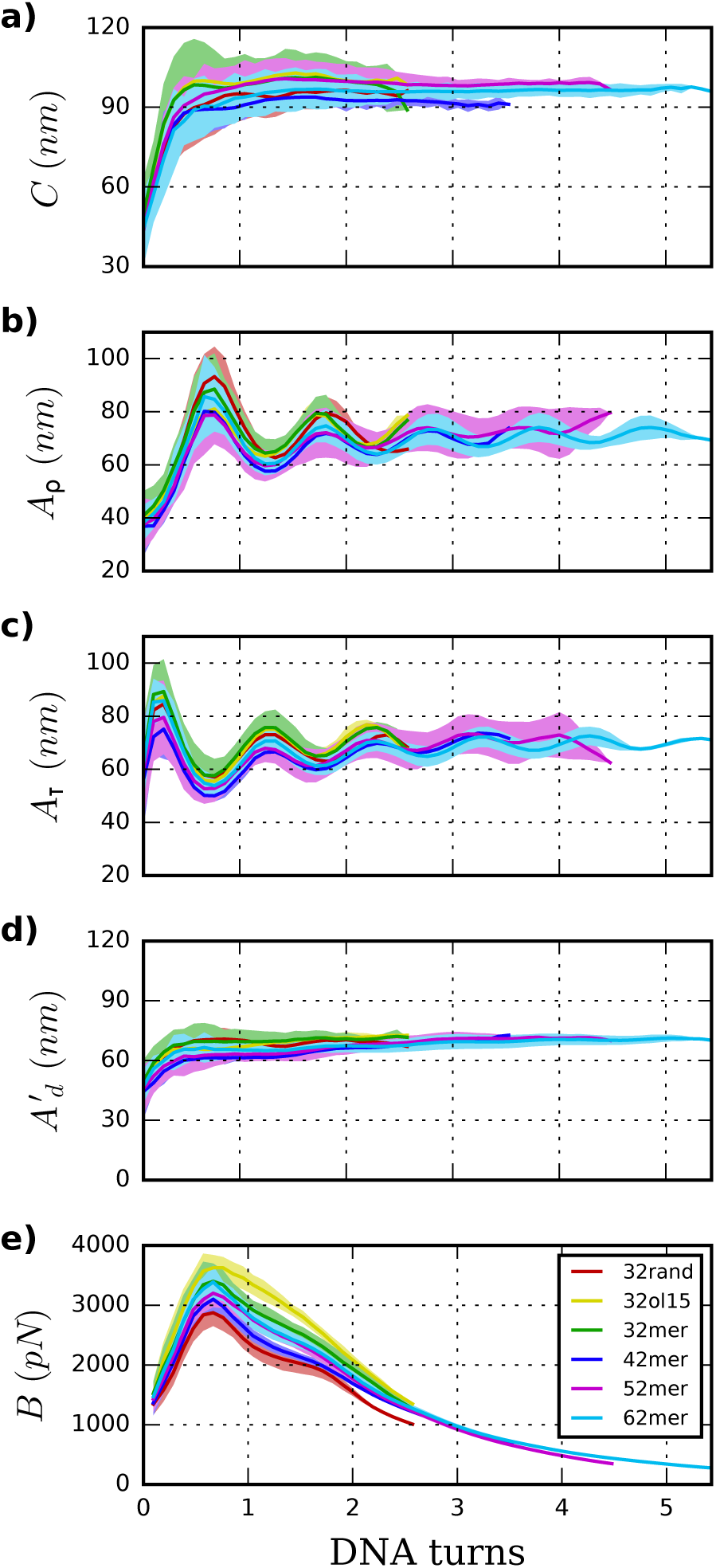
Elastic constants associated with twist (*C*), roll (*A*_*ρ*_), tilt (*A*_*τ*_) and stretch (*B*) obtained through the inverse-covariance matrix method at different lengths, together with the dynamic persistence length (*A*′_*d*_) obtained via *A*_*ρ*_ and *A*_*τ*_ combination (see text for more details). Values reported here are averages over all possible sub-fragments with a particular length, discarding the ten longest stretches, and the corresponding standard deviations are given as shade areas.

### Estimation of the long-range persistence length with its dynamic and static contributions

*SerraNA* calculates the persistence length *A* of a particular DNA fragment by means of (i) the linear fitting of the directional correlation decay or (ii) the inverse-covariance matrix method.

Mimicking the worm-like chain model (WLC), *A* is quantified by the linear approximation of the directional correlation decay between two bp tangent vectors, 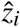 and 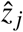 separated by an increasing number of bp steps *N* with a distance rise *b* = 0.34 nm along the DNA:^27^

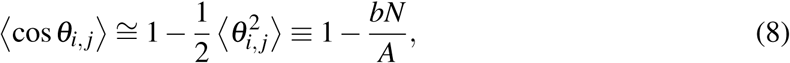

assuming a sufficiently weakly bending rod and where *N* ranges from 1 to *N∗* nucleotides, *N∗* being the longest sub-fragment considered on the fitting (see above paragraph) (Figure 4a). The static and dynamic contributions to 〈*θ*^2^〉 can be partitioned by 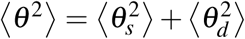, where 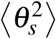 is originated from random distribution of sequence-dependent static bends and 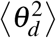 comes from the thermal fluctuations. 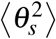 are obtained through the DNA structure rebuilt^37^ from the average base-pair step parameters. Then, the static and dynamic persistence length (*A*_*s*_, *A*_*d*_) are estimated by fitting the linear directional decay 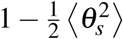 and 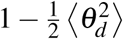, respectively (Figure 4b and 4c). *A*_*s*_ and *A*_*d*_ are combined using 1/*A* = 1*/A*_*s*_ + 1*/A*_*d*_^43^ to obtain *A* again, which should be compatible with the direct linear fit to the full bending angle correlation decay.

The inverse-covariance method provides a second estimation of the dynamic persistence length (*A*′_*d*_) by directly combining the diagonal terms of *F* corresponding to the tilt and roll elastic constants (*A*_*τ*_ and *A*_*ρ*_, respectively) for any pair of bp (Figure 3):

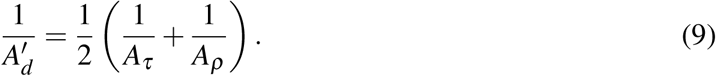

**Figure 4:**
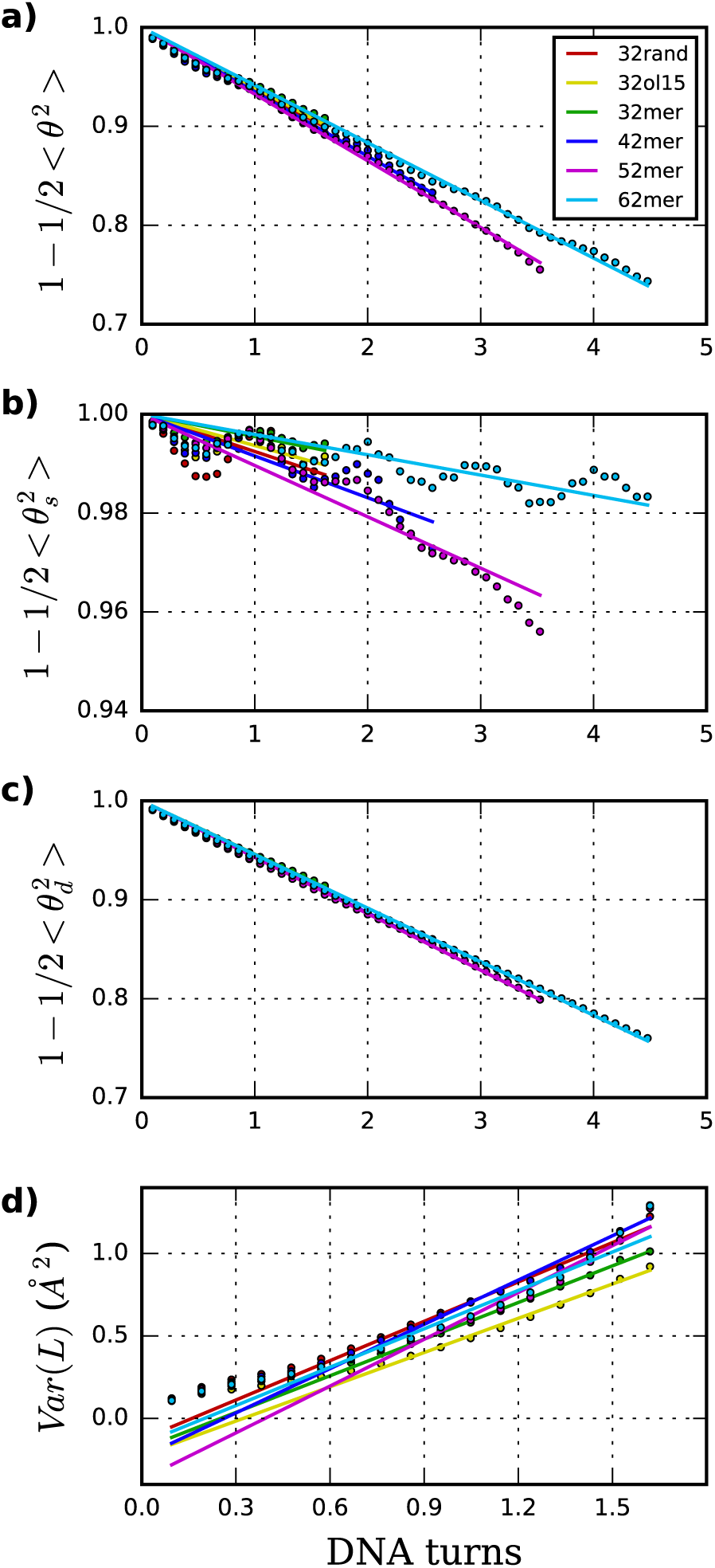
(a)-(c) Persistence length (*A*) together with its static (*A*_*s*_) and dynamic (*A*_*d*_) contributions are obtained through the linear fit of directional decays associated to 〈θ^2^〉, 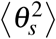 and 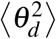, respectively. Values reported here are averages over all possible sub-fragments at a particular length, discarding ten longest lengths. (d) Stretch modulus (*B*) is obtained through linear fit of end-to-end partial variance using central 18mer.

Then, the global *A*′_*d*_ emerged from the entire DNA fragment is calculated following the methodology used for *C* (see above):

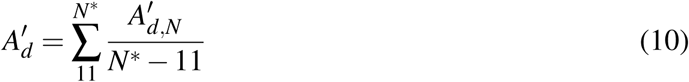

where *A*′_*d,N*_ are averages at a particular sub-fragment length with *N* bp-steps ranging from 11 to *N∗*, as the crossover from local to global dynamics occurs within the first DNA-turn (Figure 3d).

*A*′_*d*_provides higher values compared with the direct decay-fitting (*A*_*d*_) as it just considers the partial variances associated with tilt and roll (1*/Var*_*p*_(*τ*) and 1*/Var*_*p*_(*ρ*), see above) after removing their linear correlations with the other deformation variables of *F*. *A*′_*d*_is combined with the previously calculated *A*_*s*_ to obtain a second prediction of persistence length (*A′*) using the expression 1*/A*′ = 1*/A*_*s*_ + 1*/A*′_*d*_. In like manner, *A′* is stiffer than *A* as this value dismisses contributions from twist and stretch.

### Estimation of bulk stretch modulus

In a similar way to twist, stretch moduli for all subfragments *k* (*B*_*k*_) are acquired from the corresponding *F*_*k*_’s diagonal term associated to the end-to-end distance. As described before,^27^ the stretch elastic profile as a function of length presents a complex behavior due to the prevalence of stacking interactions on the shortest oligomers and the appearance of extended end-effects softening the longest DNA parts (Figure 3e and 5). In consequence, the bulk stretch modulus (*B*) is evaluated by considering only the end-to-end distances of the central 18mer and discarding oligomers shorter than 9 bp. Due to the limited number of points, the global *S* measure from a whole DNA molecule is obtained by fitting the linear increase of *Var*_*p*_(*L*) within this length range, instead of averaging the equivalent *B*_*N*_ as in the previous sections (see Figure 4d).

## Methods

### Molecular Dynamics simulations of linear DNA fragments

Linear DNA sequences of 32 bp (CGACTATCGC ATCCCGCTTA GCTATACCTA CG), 42 bp (CGCATGCATA CACACATACA TACACATACT AACACATACA CG), 52 bp (CGTATGAACG TCTATAAACG TCTATAAACG CCTATAAACG CCTATAAACG CG) and 62 bp (GCAGCAGCAC TAACGACAGC AGCAGCAGTA GCAGTAATAG AAGCAGCAGC AGCAGCAGTA GC) were extracted from the sequences 170-200 bp-long *γ*3, *γ*1, *γ*4 and *γ*2 as analyzed on Mitchell *et al.*,^26^which also correspond to the sequences NoSeq, CA, TATA and CAG on Virstedt *et al.*,^44^ respectively. DNA duplexes were built using NAB module implemented in Amber16,^45^ AMBER parm99 force-field^46^ together with parmbsc0 and parmbsc1 corrections.^36,47^ Fragments are named as 32mer, 42mer, 52mer and 62mer for the rest of the article. The 32-bp oligomer was also constructed using parmOL15^48,49^ (named 32ol15 from now on). Structures were solvated in 200 mM Na^+^ and Cl^−^ counter-ions^50^ and in TIP3P octahedral boxes^51^ with a buffer of 1.2 nm. Systems were energy-minimized, thermalized (*T* = 298 K) and equilibrated using standard protocols.^52,53^ The final structures were subject to 1 *µ*s of productive MD simulation at constant temperature (298 K) and pressure (1 atm)^54^ using periodic boundary conditions, particle mesh Ewald55 and an integration time step of 2 fs.^56^ Principal component analysis was done with pyPcazip^57^ and fast Fourier transforms were done with an in-house program written in python.

### Trajectories obtained from BIGNASim and ABC simulation databases

An extra simulation 1 *µ*s long for a DNA oligomer containing a random sequence of 32 bp (ATGGATCCAT AGACCAGAAC ATGATGTTCT CA) was obtained from the BIGNASim database^58^ and analyzed together with the above (labeled as 32random from now on). Simulation was obtained with bsc1 parameters,^36^ TIP3P water model and with neutralizing Na^+^.^50^

Elastic properties for all distinct 136 tetranucleotides were obtained by analyzing MD simulations from the ABC consortium,^23^ which are constituted of 39 oligomers of 18 bp, modeled for 1 *µ*s, using parmbsc0 force-field,^47^ SPC/E water^59^ and 150 mM K^+^Cl^−^ ion pair concentration.^60^

### MD simulation of DNA pulling

The 52 bp oligomer was stretched on a series of umbrella sampling simulations in explicit solvent following the protocol developed by Shepherd *et al.*^61^ Polymer length was increased in steps of 1 Å, which is in the range of thermal fluctuations for unconstrained DNA,^27^ thus, getting an almost instant equilibration after perturbation.^61^ DNA was pulled by a total of 8 Å, resulting in a relative extension of just approximately 5%. This early-stage stretching regime is characterized by the maintenance of all canonical interactions on the double helix (hydrogen bonding and stacking), allowing a consistent comparison with the rest of trajectories run on relaxed DNA. Each umbrella sampling window was simulated for 1 ns making a simulation 8 ns long in total.

### Linear regression and confidence intervals of elastic constants

Linear regression of directional memory and end-to-end partial variance (*Var*_*p*_(*L*)) on DNA length (*N*) are used to estimate bulk persistence lengths *A* and stretch modulus *B*, respectively, from gradients *β*_*A*_ = −*b/A* and *β*_*B*_ = *k*_B_*Tb/B*. Confidence intervals of *β*_*A*_ and *β*_*B*_ (∆*β*_*A*_ and ∆*β*_*B*_) are calculated with a confidence level of 70% in *SerraNA* using the student t-distribution for getting an almost direct comparison with other parameters where variability is estimated by standard deviation. Because *A* and *B* are non-linear functions (*y* = *f* (*x*)) of their respective gradients ∆*y* or *f* (*x* + ∆*x*) − *f* (*x*), a confidence interval can be obtained approximately by 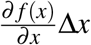. Thus, confidence intervals for *A* and *B* (∆*A* and *∆B*) are calculated by:

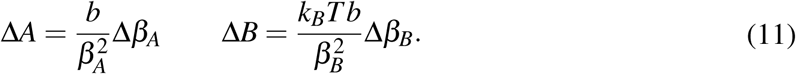

## Results and Discussion

*SerraNA* is a program written in Fortran that is freely accessible at https://github.com/agnesnoy/SerraNA under GNU Lesser General Public Licence and whose general workflow is shown in Figure 2. The program builds upon the LDEM described by Noy and Golestanian27 and it streamlines the procedure of calculating the persistence length, twist and stretch modulus of a DNA molecule or other double-stranded, helicoidal nucleic acids using an ensemble generated by MD or MC simulations.

### Torsion elastic modulus

Elastic profiles as a function of length for the whole set of simulations are presented in Figure 3. The calculated torsional modulus for all oligomers shows a crossover from the relatively soft value of around 30-60 nm at the single base-pair level to a large-scale asymptotic value between 90 and 100 nm (see Table 1), which is in agreement with previous study.^27^ While softer values at short length scales are consistent with fluorescence polarization anisotropy measurements,^62,63^ small-angle X-ray scattering (SAXS),^30^ analysis of crystallographic DNA structures^41^ and many calculations from MD,^29,36,40,64^ stiffer magnitudes concur with single-molecule experiments^8,13,14^ and other modeling estimations^20,29,65,66^ at longer length scales. Values calculated for the whole segment also fall within the long-scale range between 90-100 nm (see Table 1), achieving an overall good convergence on the microsecond-long trajectories (Figure S1)

**Table 1:** Bulk elastic constants estimated from unconstrained MD trajectories over linear DNA fragments*a*.

| DNA | $C$ (nm)<sup>b</sup> | $A$ (nm)<sup>c</sup> | $A_s$ (nm)<sup>c</sup> | $A_d$ (nm)<sup>c</sup> | $A'_d$ (nm)<sup>b</sup> | $A'$(nm)<sup>b</sup> | $B$ (pN)<sup>c</sup> |
| --- | --- | --- | --- | --- | --- | --- | --- |
| 32rand | $94.8 \pm 0.8$ | $57.4 \pm 1.6$ | $473 \pm 87$ | $65.4 \pm 0.6$ | $68.3 \pm 1.0$ | 59.7 | $1696 \pm 15$ |
| 32mer | $100.1 \pm 1.0$<br>*$101.4 \pm 1.2$* | $61.1 \pm 1.1$<br>*$58.7 \pm 1.4$*<br><u>56.3</u><br>**$50.5 \pm 2.1$** | $789 \pm 93$<br>*$562 \pm 74$* | $66.2 \pm 0.7$<br>*$65.5 \pm 0.8$* | $69.9 \pm 0.5$<br>*$69.0 \pm 1.1$* | 64.2<br>*61.4* | $1920 \pm 18$<br>*$2207 \pm 41$* |
| 42mer | $92.8 \pm 0.7$ | $54.8 \pm 0.6$<br><u>54.8</u><br>**$45.5 \pm 0.5$** | $422 \pm 24$ | $63.0 \pm 0.6$ | $64.7 \pm 2.3$ | 56.1 | $1705 \pm 12$ |
| 52mer | $99.2 \pm 0.8$ | $52.9 \pm 0.2$<br><u>51.5</u><br>**$45.5 \pm 0.8$** | $344 \pm 10$ | $62.6 \pm 0.2$ | $67.8 \pm 2.9$ | 56.6 | $1843 \pm 27$ |
| 62mer | $96.1 \pm 0.6$ | $61.2 \pm 0.3$<br><u>51.5</u><br>**$41.7 \pm 0.5$** | $869 \pm 36$ | $65.8 \pm 0.2$ | $68.2 \pm 1.8$ | 63.3 | $1731 \pm 9$ |
| Average | $96.6 \pm 2.7$ | $57.5 \pm 3.3$ | $579 \pm 210$ | $64.6 \pm 1.5$ | $67.8 \pm 1.7$ | $60.0 \pm 3.3$ | $1779 \pm 88$ |

<sup>a</sup> Elastic constants obtained using OL15 force-field are in italics. Persistence lengths on sequences over 100 bp, from which short fragments has been extracted from (see Methods), are in underlined text when they come from MC simulations<sup>26</sup> and in bold when they come from experiments.<sup>44</sup>
<sup>b</sup> Overall averages and standard deviations for elastic constants obtained through the inverse-covariance method (twist $C$ and persistence length $A'$ and $A'_d$ ) are calculated using the different sub-fragment lengths between 11 bp-steps to $N^*$ , $N^*$ being the maximum number of bp considered (see Methods).
<sup>c</sup> Overall values and confidence levels at 70% for persistence lengths following the WLC model ( $A$ , $A_s$ and $A_d$ ) and stretch modulus ( $B$ ) are obtained through linear fits (see Methods).

### Persistence length

Persistence length (*A*), as well as its static (*A*_*s*_) and dynamic (*A*_*d*_) components, were deduced following the principles of the WLC model. Persistence lengths calculated by the fitting of directional decays are in general higher than the corresponding experimental data^44^ and coarse-grained modeling^26^ (see Table 1), although it should be noted that our magnitudes are obtained with much shorter DNA molecules. Our average across sequences gives an overall stiffer estimation (57± s.d. 3 nm) compared with the range of experimental measurements (45 − 55 nm)^6–10^ but in general agreement with estimations from simulations.^26,29,65^ Part of this difference might be originated from the fact that our simulations are obtained with fully controlled ionic solutions (200mM NaCl), without containing Mg^2+ 44^ and other buffers like Hepes, Tris or EDTA^7–10^ known to affect DNA flexibility.^6,29,67^ This variation could also be caused by inaccuracies in the modeling methods, although it is difficult to assess without comparing exactly the same sequences and with such a limited number of oligomers.

Figure 4 shows tangent-tangent correlations arisen from *A*_*s*_ (*ie* from intrinsic curvature) exhibit modulations in phase with DNA-turn periodicity, in contrast to the decay originated from *A*_*d*_ (*ie* from thermal fluctuation).^27^ Our calculations indicate *A*_*s*_ is much stiffer than *A*_*d*_, even though *A*_*s*_ is the main source of variability (*A*_*s*_ = 576 ± 191 nm; *A*_*d*_ = 64.7 ± 1.4 nm; see Table 1 and Figure 4). This trend was already observed on MC simulations^26^ and it would explain the difficulty of arriving to a consensus description by experiments (*A*_*s*_ > 1000 and *A*_*d*_ ≈ 50 nm;^68^ *A*_*s*_ ≈ 130 and *A*_*d*_ ≈ 80 nm^69^). For atomistic simulations, the small and oscillating decay together with the limited molecular length make the estimation of *A*_*s*_ (and as a consequence *A*) challenging and sometimes imprecise. These sources of error are exposed by the broad confidence intervals of *A*_*s*_ compared to *A*_*d*_ (Table 1) and the relative lack of convergence in some of *A*_*s*_ measurements e.g. for the 62mer (Figure S1). Another example is the discrepancy of *A* obtained by two different DNA force-fields (BSC1 and OL15), which is mainly caused by *A*_*s*_ and not *A*_*d*_ (see Table 1), being complicated to judge whether error comes from force-fields or the linear fit.

The inverse-covariance method yields an increased dynamic persistence length *A*′_*d*_of 68 nm, and a resulting persistence length *A′* of 60 nm, as it only considers fluctuations not correlated with other deformation variables (*ie* partial variances, see methods). *A*′_*d*_is calculated through the combination of roll and tilt elastic constants, *A*_*ρ*_and *A*_*τ*_, which produce periodic and antisymmetric profiles as a function of fragment length due to bending anisotropy towards grooves and backbone (see Figure 3). On lengths containing half and complete helical turns, *A*_*ρ*_ and *A*_*τ*_ are equivalent because grooves and backbone face equitably towards both bending axes, whereas, at intermediate lengths, there is an imbalance between them (see Figure 1).

### Stretch Modulus

Stretch modulus deduced from all unconstrained simulations present a non-monotonic dependence on length similar to the one previously described by Noy and Golestanian^27^ and reproduced by Wales and co-workers^29^ (see Figure 3e). Base-stacking interactions cause stiffening at short scales up to 7 bp length as elastic constants present similar values associated with contour-lengths (see Figure S2). For longer sub-fragments, cooperativity emerges due to coordinated motion, softening the stretch modulus in two stages: (i) towards a plateau that would correspond to the regime captured by force-extension experiments^11,12^ after incorporating an internal mode 13 bp long^27^ and (ii) towards much more flexible magnitudes originated by long-ranged end-effects.^27^ Principal component analysis reveals a mode that essentially captures vibration from edges and that produces a proportionate influence over the different oligomers containing gradually more bp (see Figure 5). This fact shows that the characteristic length of the stretching-end mode is longer than five DNA-turns, still not reached for our atomistic simulations. In contrast, *L* increments are uniformly distributed along the molecule in the simulation where DNA is actively pulled (see Figure 5), which shows that the end-stretching motion is just a vibrational mode not relevant for extracting the intrinsic stretch modulus of DNA.

**Figure 5:**
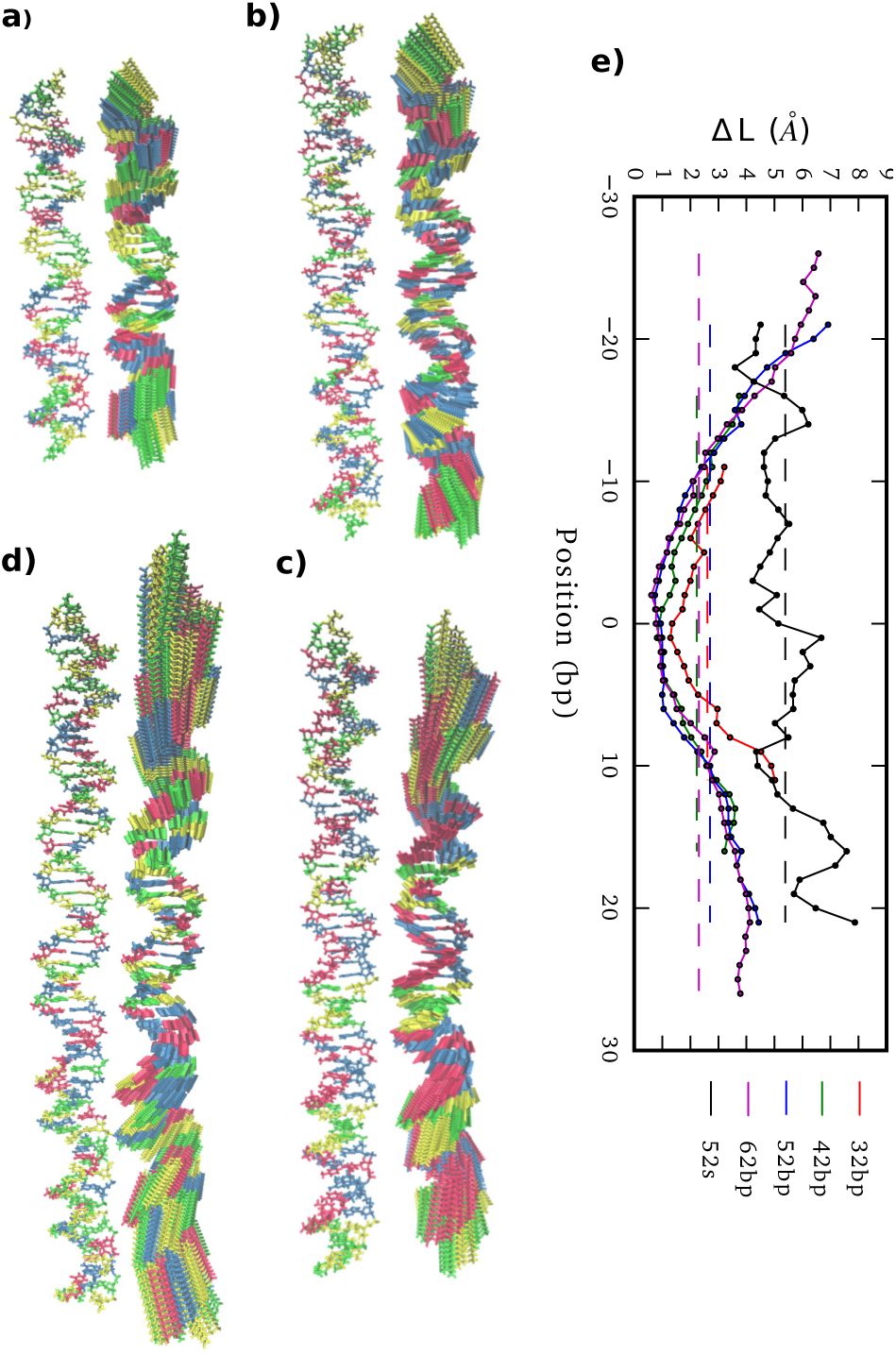
(a)-(d) 32mer, 42mer, 52mer and 62mer averaged structures together with the corresponding end-effect essential modes. (e) Molecular position dependence of end-to-end distance local increments (∆*L*) caused by end-vibrational modes from relaxed simulations and by the 52 bp pulling simulation with a maximum extension of 5% (52s), using in all cases sub-fragments of 5 bp length.

We estimate stretch modulus via linear fitting of *Var*_*p*_(*L*) just using the central 18 bp, since they constitute the molecular domain significantly unaltered by end-effects (see Figure 5). Results give an overall average of 1779 ± 88 pN (Table 1), which is reasonably close to the experimental value ca. 1500 pN.^12^

### From local to global elastic behavior

By analyzing elastic and structural length-dependence, *SerraNA* can also reveal how global elastic constants build up from the dynamics of smaller scales. For example, Figure 6 compares the length-evolution on bending angles of the more bendable fragment (52mer) with the less one (62mer). Interestingly, bending is comparable between the two sequences at the single bp-step level (7.1±1.5 and 7.2±1.1 degrees, respectively), but are able to cause distinct values at the longer scale of 38 bp (35.6±1.6 and 33.0± 0.7 degrees). The main difference at intermediate lengths (8, 16 and 28 bp) is the higher degree of periodicity, which is in phase with DNA helicoidal shape, presented by the curved oligomer compared to the straight one (see Figure 6). Our data suggests that for creating a regular pattern characteristic of the curved fragment, a frequency with an exact number of cycles per DNA-turn at the single bp-step level is needed (3 cycles per DNA-turn for the 52mer in front of 3.5 for the 62mer, see Figure 6), so local bends can couple for building up a significant curvature. Our results are in the same line of others that highlighted the importance of periodicity^32^ for understanding the special mechanical properties of A-tracts^19^ or nucleosome-positioning sequences.^70,71^

**Figure 6:**
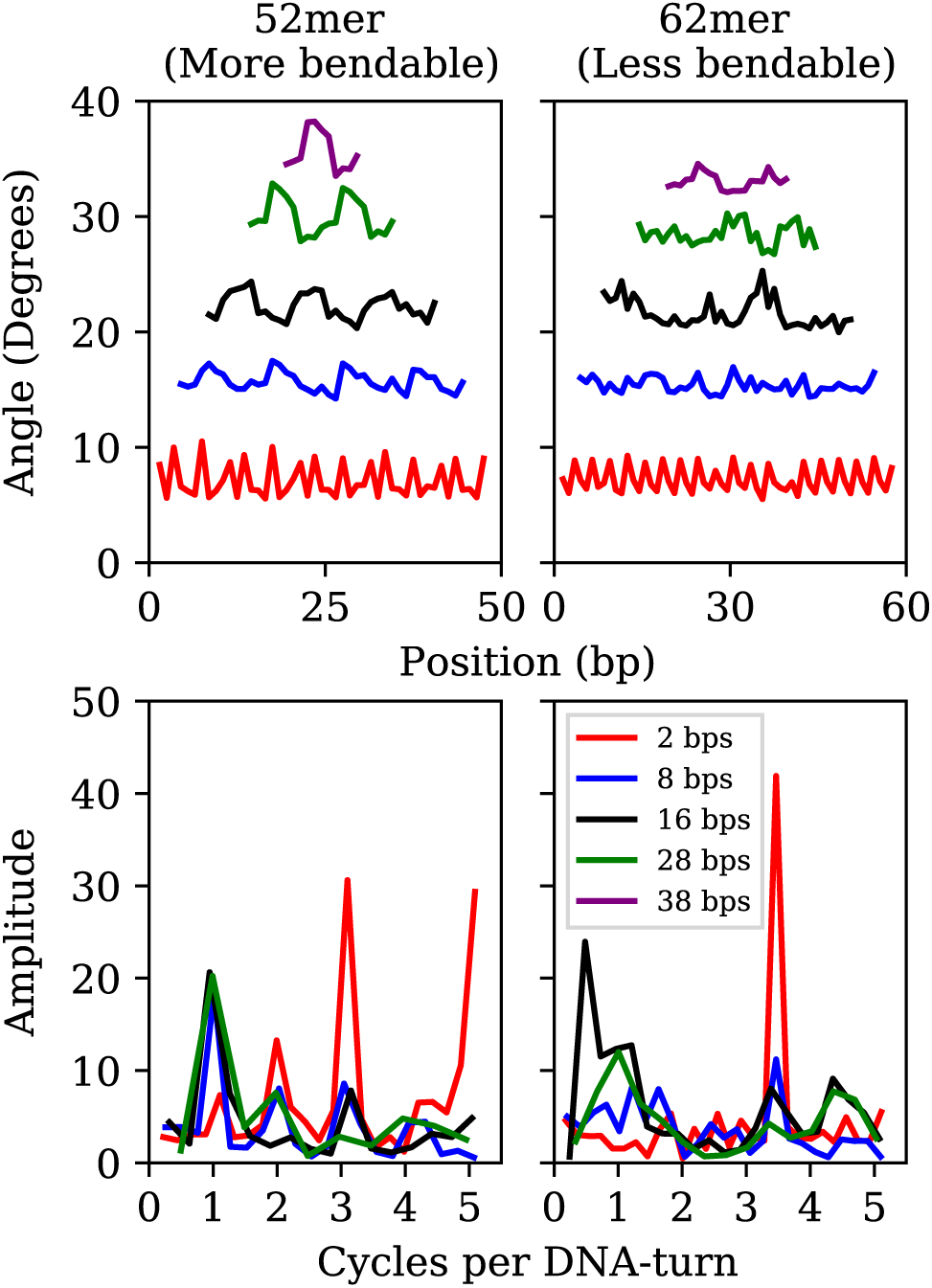
Top: length-evolution of bending angle profiles along the sequence for the most (left) and less (right) curved oligomers (52mer and 62mer, respectively). Bottom: frequencies (in cycles per DNA-turn) obtained after applying fast Fourier transforms to bending positional data

### Tetranucleotide elastic constants from ABC database

Lastly, we analyzed how different DNA elastic properties depend on sequence. To this end, we applied *SerraNA* to the ABC simulation database, which contains the whole set of 136 tetra-nucleotide sequences in 39 different oligomers^23^ (see Figure 7). In general, we can observe a high degree of variability with flexible sequences twice as soft as rigid ones for all elastic constants.

**Figure 7:**
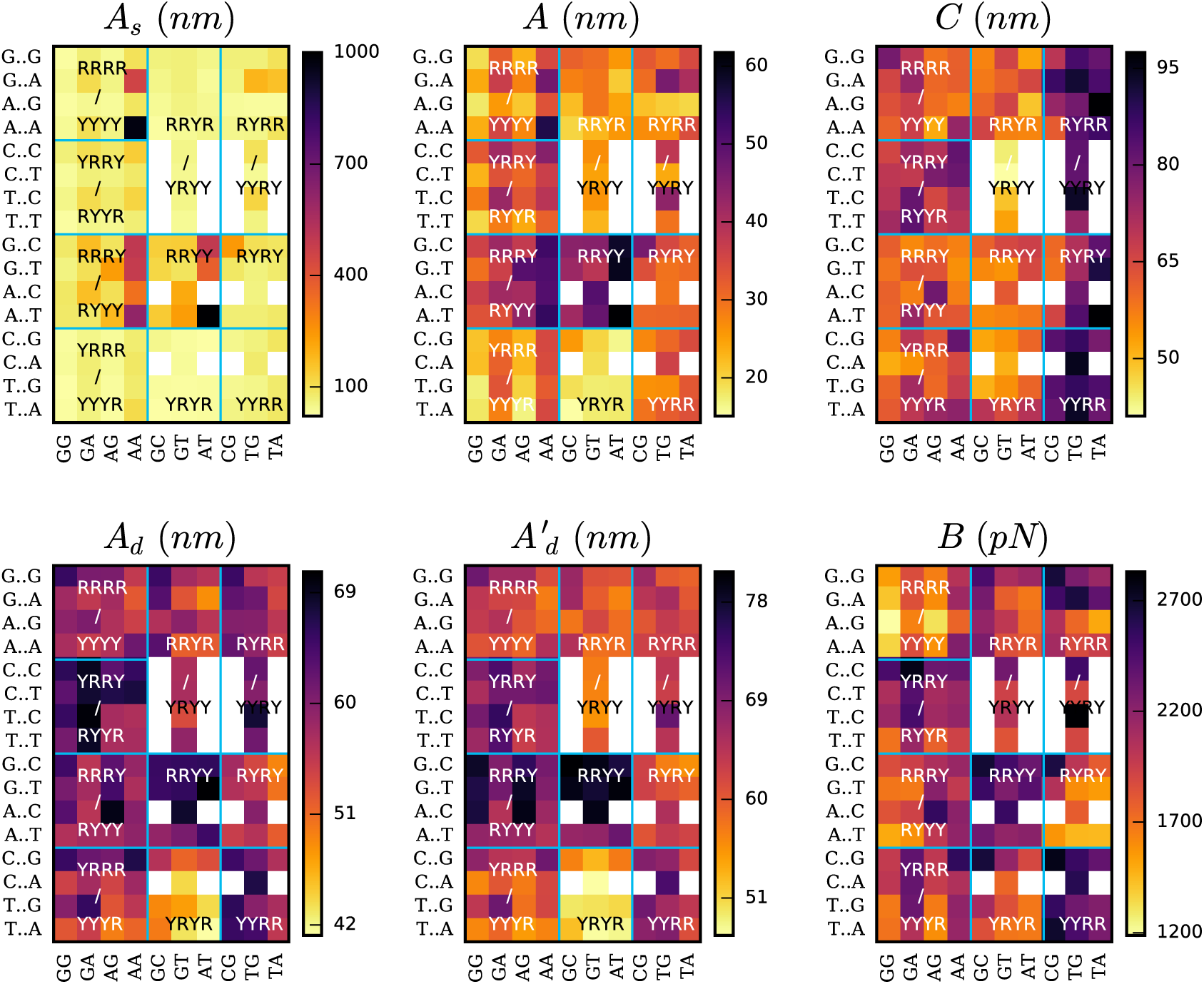
Elastic constants at the length of 4 bp for the whole set of 136 tetra-nucleotide sequences obtained from ABC simulation database. Total persistence length together with its static and dynamic components (*A*, *A*_*s*_ and *A*_*d*_, respectively) are calculated using the directional decay at the tetramer length. Twist (*C*), stretch modulus (*B*) and the second estimation of dynamic persistence length (*A*′_*d*_) are obtained directly from the inverse-covariance matrix for tetranucleotides. Vertical axis indicates middle steps, and horizontal axis flanking bases. Horizontal and vertical lines organize sequences according purine (R) or pyrimidine (Y) type. Sequence duplication is excluded through the use of white squares. AATT’s *A*_*s*_ is off the palette with a value of 1267±144 nm (see Table S6).

The static persistence length is the most variable parameter in sequence space, spanning almost two orders of magnitude: from <25 nm in the case of TGGG, TGCA and CATG to >1000 nm for AATT (see Tables S1-S10). In general we observe that the majority of the tetramers are very flexible and just 13 sequences (9%) have values >200 nm. The less curved tetramers involve central AA or AT steps, with AATT and AAAA being the top two with 1267 and 970 nm, respectively (Figure 7 and Table S1 and S6). This is in agreement with previous studies and with the idea of A-tracts being so stiff that they impair nucleosomes wrapping^72^ but facilitate looping and gene regulation when they are placed in phase.^72–74^ It’s worth mentioning that the extremely low values presented by most of the sequences are characteristic of this particular length (4 bp) as there is an accumulation of bending towards the major groove on one DNA side.^32^ This behavior is reflected in the oscillations of the directional curvature correlation^27^ (see Figure 4b) and is exploited in fundamental processes like protein:DNA recognition^32,75^ and the formation of DNA loops.^76^

When looking at the effect of thermal fluctuations on bendability (Figure 7), we recover a scenario in agreement with previous crystallographic and modeling studies^21,23,64^ where sequences containing the maximum number of “hinges” YR bp-steps (YRYR) are the most flexible and sequences with just RR and RY steps (RRYY and RRRY) the most rigid (being Y pyrimidines and R purines). Within the last two types of tetramers, sequences presenting central AA or AT steps are especially stiff (55±4 nm, see Table S3 and S6) due to the influence of curvature, whereas YRYR tetramers containing TA and CA are specially flexible (17±1 nm, see Table S7).

The two different estimations of dynamic persistence lengths provide similar patterns on the sequence space in spite of their different ranges. We observed that although static persistence length presents more disparate values than the dynamic component, the latter is also important in determining the relative elasticity across all oligomers.

Figure 7 shows that torsional moduli ranges from approximately 40 nm, which are characteristic of dinucleotides, up to 90 nm, which is a value typical of long scales. This is because 4 bp constitute an intermediate length in the transition from local to bulk behavior (see Figure 3), so the levels of correlation between bp-step fluctuations tend to diverge (see Figure 2c), making sequence-dependence analysis very convoluted. Broadly, the most rigid sequences for this parameter are the ones with a central YR step (Figure 7), which strikingly is the most flexible bp-step type at the dinucleotide level (see Figure S3) in agreement with previous studies.^21,52,64^ We also observe that sequences with a bimodal behavior in the central step^22^ don’t show any special flexible feature. These facts demonstrate the remarkable importance of flanking bases in building up overall fluctuations and the very complex interplay between dinucleotide steps.^77,78^

Stretch modulus at the tetra-bp length are relatively high (see Figure 7 and Table S1-S10) compared with experiments at the long-range scale. The remarkable similarity between other distance definitions (*ie* end-to-end, contour length or added-rise, Figure 7 and S4) suggests stretch stiffness at this length is mainly influenced by the strong stacking interactions. There is also an important degree of variability among sequences with some steps like AGGG, AGGA and AAGG presenting stretch modulus <1400 pN, which are twice as flexible as others such as CGAC, TTGC and CCGG (>2700 pN). In general, we observe YYRR and RRYY steps be the most rigid and RRRR the most flexible for this parameter, being determined mainly by the vertical component (added-rise) but also with some influence from lateral displacements, in particular from slide direction. AAAA sequence is an exception of RRRR type of tetramer by presenting a relatively stiff stretch modulus (2241±88, see Table S1), in reasonably good agreement with recent experimental data (∼2400 pN)^19^

The analysis of ABC database makes clear that there is a flexibility dependence on the sequence of DNA and that reasonably extends to sequences larger than 4 bp. Regarding tetranucleotides elastic constants, rigidity tends to increase in regions composed by RRYY, YYRR, RRRY and YRRY, whereas sequences made with YRYR, RRYR are in general flexible, although this classification strongly depends on the type of elastic parameter.

## Concluding Remarks

In this article we present *SerraNA*, which is an open code that describes the elastic properties of nucleic-acids molecules with a canonical helicoidal shape (B- or A-form) using ensembles obtained from numerical simulations. We apply the program to analyze a series of atomistic MD simulations over DNA fragments and compare the extracted elastic values with available experiments.

We find reasonably good agreement on stretch and torsional modulus between our estimations (97±3 nm and 1778±88 pN) and experimental values (around 100 nm and 1500 pN, respectively). The calculation of stretch modulus is especially challenging because of the end-stretching vibration that masks the thermal fluctuations characteristic of the experimental stretch modulus at the range of kbp. As atomistic simulations are done over relatively short DNA molecules (tens to a hundred of bp), *SerraNA* approximates the calculation of this elastic parameter using only the two central DNA turns. In spite of all approximations, we find remarkable agreement between the only sequence experimentally measured, the A-tract, (∼2400 pN)^19^ and the modeled AAAA tetramer (2241±88).

In the case of persistence length, simulations provide a slightly more rigid measure (57±3 nm) than the generally accepted value of 50 nm, although it’s hard to discern whether it is due to intrinsic problems of force-fields, to non-identical ionic conditions with experimental buffers or to trouble in measuring the static persistence. Modulations on the tangent-tangent decay caused by DNA intrinsic shape and the relative shortness of the simulated DNA fragments makes the calculation of the static component of persistence length peculiarly complicated. Moreover, our simulations indicate that DNA curvature is the main source of variability on bendability between sequences (510±210 nm), compared with 64.6±1.5 nm caused purely by thermal fluctuations. When we analyze the whole set of tetra-bases sequences from ABC database, we observed again a higher degree in variation on the static persistence length (s.d. 159 nm) in contrast to dynamic persistence length (s.d. 5.9 nm) (see Table S11).

*SerraNA* also indicates how global elasticity emerges from local fluctuations by analyzing the change of mechanical properties as the length of considered fragments is systematically increased. Because the crossover from single base-pair level to bulk elastic behavior occurs typically within one helical turn of DNA, relatively short DNA fragments, like the ones simulated here, are already useful for uncovering this effect. In the case of persistence length, our results show that periodic patterns in phase of the DNA helical turn are particularly advantageous for developing significant bendability at longer scales.

Finally, the systematic analysis of the whole set of 136 tetranucleotides reveals big differences with some sequences doubling others in all elastic parameters and, as a consequence, indicates the importance of sequence in determining DNA elastic properties. YRYR are the most flexible sequences compared with RRYY and RRRY, which are the most rigid. Particularly, AT and AA are the bp-steps causing less bendability, due to its straight natural configuration, in contrast to the highly flexible TA and CA bp-steps. RRYY and RRRY tetramers containing AT and AA steps present a persistence length 38 nm higher than YRYR tetramers with TA and CA steps. This demonstrate the role of AT-rich motifs in defining opposite mechanical properties, which can build up global deformability on longer sequences when they are regularly phased with the helicoidal shape. We thus see that *SerraNA* can shed light on the reasons behind the different emerging mechanical properties between AT and GC-rich long sequences^16,19^ and, consequently, how their different biological functions might occur.^79^

In general, thought, we observe a complicated dependence for the different type of tetra steps compared with the dinucleotide level, showing the relevance of flanking sequences and the complex interplay between the different bp-steps. We expect that the use of *SerraNA* will help to clarify further how DNA elasticity can be modulated as a function of sequence, having important implications in understanding fundamental processes like DNA-protein recognition, DNA looping or packing inside the cell.^80^ In particular, we anticipate using SerraNA in a range future experimental investigations^81^ which will help us to unravel new physical properties of DNA at the single-molecule level.^82,83^

## Acknowledgement

We thank John Maddocks, Richard Lavery, Modesto Orozco and the rest of ABC consortium for giving us accessibility to all simulations of the database. This work was supported by the EPSRC grant EP/N027639/1 and by the Leverhulme Trust grants RPG-2019-156 and RPG-2017-340. V.V-B. was funded by CONACYT agency from Mexican government (scholarship no 291163) and M.B. by EPSRC (EP/R513386/1). Computational time was secured on ARCHER and JADE via the UK High-End Computing Consortium for Biomolecular Simulation, HECBioSim, supported by EPSRC grant EP/R029407/1 and on Cambridge Tier-2 system funded by EPSRC Tier-2 capital grant EP/P020259/1. We also thank Tier 3 High Performance Computing (HPC) facilities at York (Viking and YARCC clusters) for additional computational resources.

