## Supplementaty figures for "*SerraNA*: a program to determine nucleic acids elasticity from simulation data"

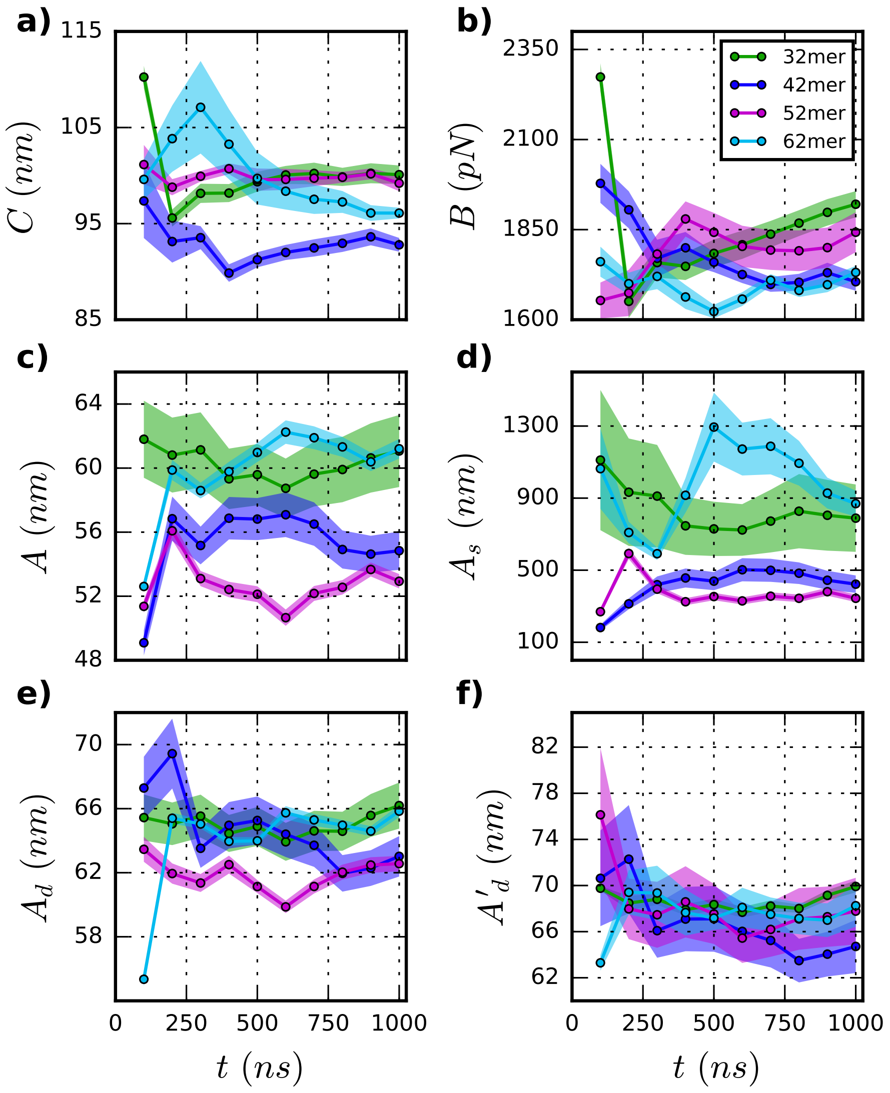


Figure 1: Elastic constants for twist $C$, stretch $B$, persistence length $A$, static persistence length $A_s$, dynamic persistence length $A_d$ and a second prediction of the dynamic persistence length $A'_d$ for increasing extensions (in ns) of the trajectories over DNA fragments containing 32 (green), 42 (blue), 52 (purple) and 62 (cyan) bp long. Shaded areas represent standard deviations in the case of $C$ and $A'_d$, and uncertainty values with 70% of confidence level for variables obtained through linear fits ($B$,$A$,$A_s$ and $A_d$).

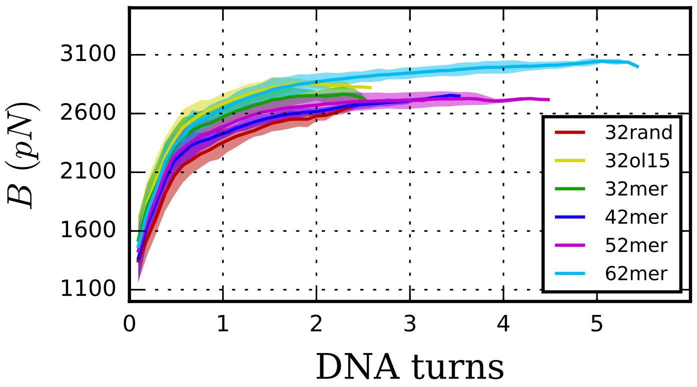


Figure 2: Stretch Modulus $B$ at different lenghts using contour length instead of end-to-end distance and including simulations for a 32bp random sequence (32rand) and the 32mer using parmOL15 force field (32ol15) (see Methods).

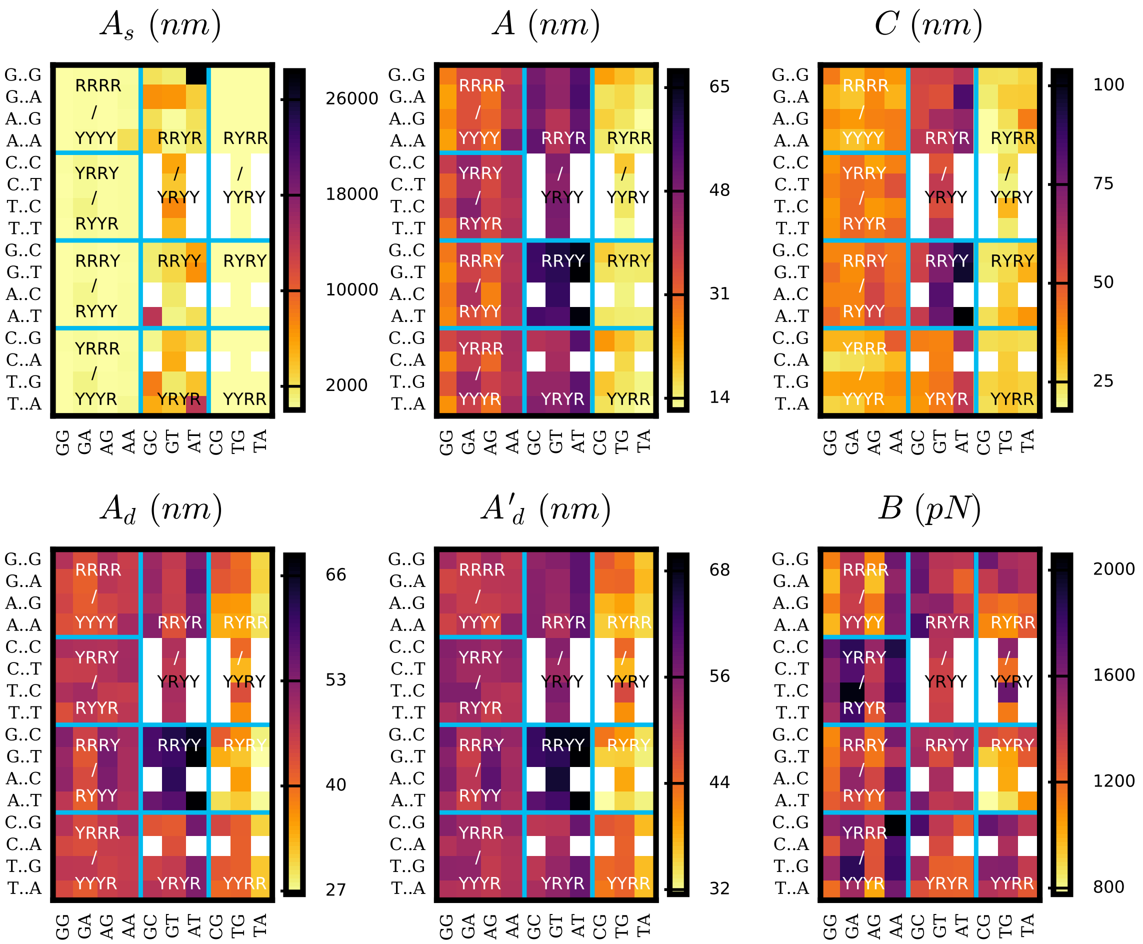


Figure 3: Elastic constants at the dinucleotide length for the whole set of 136 tetra-nucleotide sequences from ABC simulation database. Total persistence length together with its static and dynamic components ($A$, $A_s$ and $A_d$, respectively) are calculated using the directional decay at dimer level. Twist ($C$), stretch modulus ($B$) and the second estimation of dynamic persistence length ($A'_d$) are obtained directly from the inverse-covariance matrix for dinucleotides. Vertical axis indicates middle steps, and horizontal axis flanking bases. Horizontal and vertical lines organize sequences according purine (R) or pyrimidine (Y) type. Sequence duplication is excluded through the use of white squares.

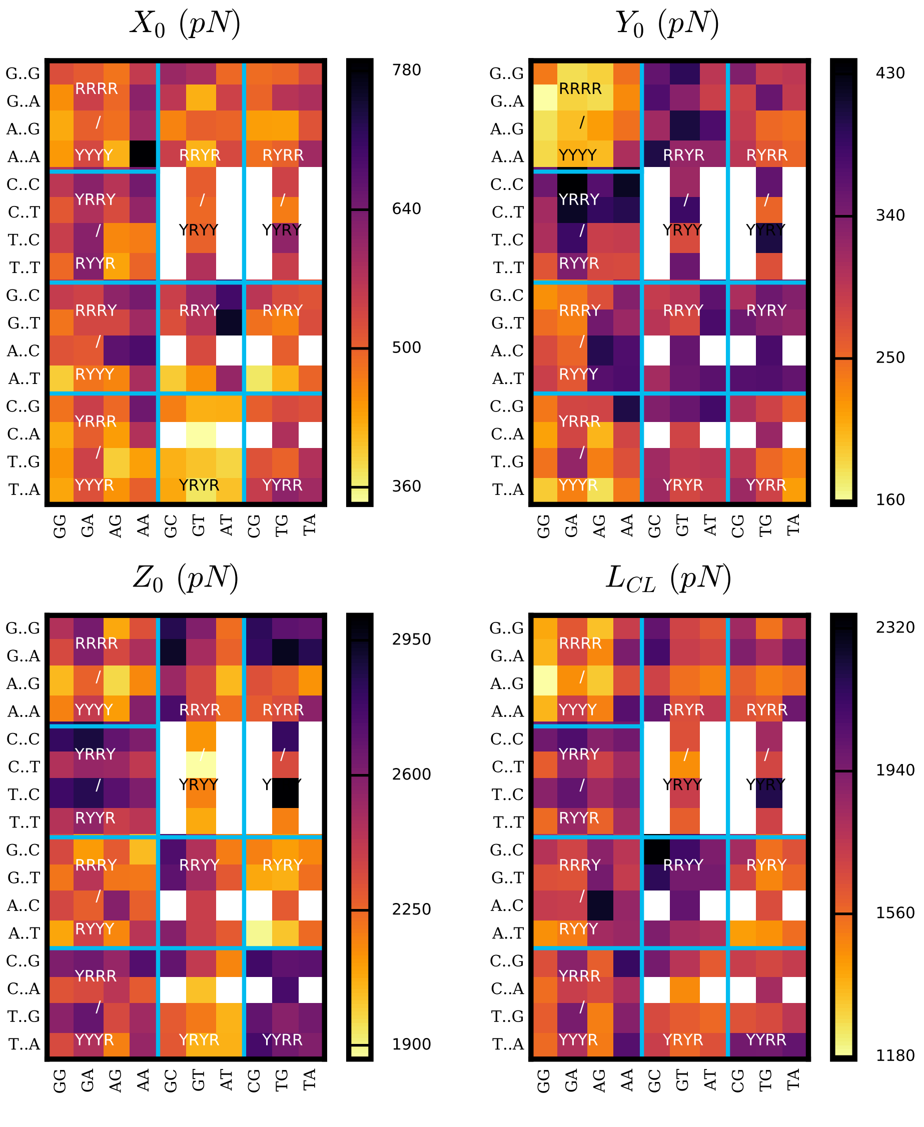


Figure 4: Added shift $X_0$, slide $Y_0$, rise $Z_0$ and contour length $L_{CL}$ elastic constants at the 4 bp length for the 136 tetranucleotide sequences from ABC simlation databse. Vertical axis represent the flanking base-pairs and the horizontal axis represents the middle base-steps. Horizontal and vertical lines organize sequences according purine (R) or pyrimidine (Y) type. Sequence duplication is excluded through the use of white squares.

Table 1: Persistence length $A$, its static $A_s$ and dynamic $A_d$ components and twist $C$ and stretch modulus $B$ of RRRR tetranucleotide sequences.

| Sequence | $A$ $(nm)$ | $A_d$ $(nm)$ | $A_s$ $(nm)$ | $C$ $(nm)$ | $B$ $(pN)$ |
|---|---|---|---|---|---|
| GGGG | 19.0 ± 0.6 | 65.4 ± 2.6 | 26.9 ± 1.1 | 79.1 ± 6.4 | 1541 ± 47 |
| GGGA | 24.0 ± 0.7 | 57.9 ± 1.1 | 41.0 ± 2.2 | 65.8 ± 2.0 | 1411 ± 34 |
| AGGG | 18.0 ± 0.5 | 59.3 ± 1.7 | 25.8 ± 0.7 | 63.7 ± 3.1 | 1188 ± 26 |
| AGGA | 23.5 ± 2.2 | 57.2 ± 0.8 | 40.3 ± 6.3 | 61.0 ± 0.8 | 1358 ± 39 |
| GGAG | 31.9 ± 0.6 | 60.7 ± 1.1 | 67.6 ± 3.9 | 69.3 ± 2.1 | 1863 ± 6 |
| GGAA | 36.7 ± 0.6 | 55.5 ± 0.7 | 108.1 ± 2.7 | 74.5 ± 4.9 | 1844 ± 44 |
| AGAG | 25.5 ± 2.1 | 60.5 ± 1.0 | 44.5 ± 6.4 | 66.9 ± 4.8 | 1637 ± 78 |
| AGAA | 35.4 ± 5.0 | 55.0 ± 4.3 | 101.8 ± 26.9 | 66.7 ± 1.7 | 1863 ± 136 |
| GAGG | 23.3 ± 1.6 | 60.6 ± 0.9 | 37.9 ± 3.8 | 58.6 ± 1.6 | 1652 ± 122 |
| GAGA | 27.7 ± 0.8 | 57.6 ± 0.5 | 53.4 ± 3.0 | 60.1 ± 1.7 | 1603 ± 38 |
| AAGG | 21.2 ± 1.0 | 56.8 ± 0.5 | 34.0 ± 2.7 | 59.2 ± 4.5 | 1311 ± 23 |
| AAGA | 27.4 ± 4.7 | 56.6 ± 2.6 | 54.7 ± 14.7 | 51.8 ± 8.4 | 1581 ± 84 |
| GAAG | 34.9 ± 1.3 | 55.4 ± 0.5 | 95.6 ± 10.6 | 61.3 ± 1.4 | 2028 ± 109 |
| GAAA | 46.9 ± 0.1 | 52.3 ± 0.1 | 454.1 ± 20.1 | 61.5 ± 2.0 | 2127 ± 24 |
| AAAG | 33.7 ± 0.4 | 54.6 ± 1.1 | 87.8 ± 0.1 | 56.8 ± 3.0 | 1754 ± 47 |
| AAAA | 55.8 ± 7.0 | 61.6 ± 3.4 | 970.0 ± 506.1 | 74.6 ± 6.6 | 2241 ± 88 |
| Average | 30.3 ± 9.9 | 57.9 ± 3.2 | 140.2 ± 236.0 | 64.4 ± 7.0 | 1688 ± 288 |

Table 2: Persistence length $A$, its static $A_s$ and dynamic $A_d$ components and twist $C$ and stretch modulus $B$ of YRRY tetranucleotide sequences.

| Sequence | $A\ (nm)$ | $A_d\ (nm)$ | $A_s\ (nm)$ | $C\ (nm)$ | $B\ (pN)$ |
|---|---|---|---|---|---|
| TGGT | 19.6 ± 0.8 | 61.1 ± 3.8 | 28.8 ± 1.0 | 72.9 ± 5.4 | 1700± 109 |
| TGGC | 28.4 ± 2.2 | 65.5 ± 2.4 | 50.4 ± 5.4 | 69.9 ± 2.1 | 2139± 82 |
| CGGT | 23.0 ± 0.5 | 62.7 ± 2.5 | 36.4 ± 0.6 | 69.0 ± 4.5 | 2072± 88 |
| CGGC | 32.7 ± 0.6 | 66.2 ± 2.3 | 64.9 ± 2.6 | 65.8 ± 2.7 | 2468± 82 |
| TGAT | 28.5 ± 1.6 | 68.7 ± 3.4 | 48.8 ± 3.2 | 82.3 ± 2.5 | 2153± 118 |
| TGAC | 42.3 ± 1.7 | 70.1 ± 1.9 | 106.7 ± 6.6 | 81.0 ± 0.4 | 2470± 39 |
| CGAT | 33.6 ± 0.7 | 68.1 ± 0.7 | 66.5 ± 2.9 | 68.3 ± 2.8 | 2421± 44 |
| CGAC | 41.5 ± 1.2 | 69.0 ± 2.7 | 104.2 ± 1.7 | 74.1 ± 1.4 | 2754± 65 |
| TAGT | 24.0 ± 1.9 | 57.2 ± 2.8 | 41.4 ± 4.3 | 68.0 ± 4.9 | 1771± 62 |
| TAGC | 33.1 ± 0.8 | 57.4 ± 0.9 | 78.1 ± 3.7 | 65.4 ± 1.9 | 2119± 49 |
| CAGT | 30.9 ± 1.1 | 66.9 ± 1.8 | 57.6 ± 3.2 | 78.2 ± 1.8 | 2097± 26 |
| CAGC | 35.5 ± 1.7 | 62.8 ± 2.4 | 81.8 ± 5.4 | 70.2 ± 2.0 | 2329± 70 |
| TAAT | 31.6 ± 2.5 | 57.4 ± 2.3 | 70.7 ± 9.0 | 69.5 ± 2.0 | 1923± 68 |
| TAAC | 38.8 ± 3.6 | 56.8 ± 2.4 | 124.0 ± 23.3 | 72.0 ± 2.6 | 2171± 16 |
| CAAT | 36.6 ± 2.5 | 67.4 ± 2.9 | 80.2 ± 7.9 | 80.5 ± 2.4 | 2147± 81 |
| CAAC | 44.6 ± 1.6 | 65.5 ± 2.4 | 139.7 ± 5.9 | 81.5 ± 0.9 | 2347± 125 |
| Average | 32.8 ± 6.9 | 63.9 ± 4.5 | 73.8 ± 30.7 | 73.0 ± 5.7 | 2193± 261 |

Table 3: Persistence length $A$, its static $A_s$ and dynamic $A_d$ components and twist $C$ and stretch modulus $B$ of RRRY tetranucleotide sequences.

| Sequence | $A\ (nm)$ | $A_d\ (nm)$ | $A_s\ (nm)$ | $C\ (nm)$ | $B\ (pN)$ |
|---|---|---|---|---|---|
| GGGT | 29.7 ± 1.3 | 60.8 ± 0.2 | 58.5 ± 5.3 | 58.4 ± 1.2 | 1682 ± 113 |
| GGGC | 35.9 ± 2.2 | 64.9 ± 2.2 | 80.5 ± 7.8 | 61.5 ± 2.0 | 1861 ± 105 |
| AGGT | 33.5 ± 1.7 | 56.7 ± 2.0 | 82.3 ± 7.8 | 60.5 ± 0.1 | 1498 ± 82 |
| AGGC | 36.5 ± 0.7 | 63.3 ± 1.4 | 86.5 ± 4.7 | 61.0 ± 3.0 | 1833 ± 71 |
| GGAT | 36.4 ± 0.6 | 57.6 ± 0.4 | 99.1 ± 5.8 | 63.9 ± 0.7 | 1670 ± 69 |
| GGAC | 41.9 ± 2.1 | 56.0 ± 0.9 | 169.0 ± 25.4 | 55.8 ± 0.0 | 1938 ± 46 |
| AGAT | 33.7 ± 0.4 | 57.6 ± 0.4 | 81.5 ± 3.3 | 74.0 ± 1.8 | 1589 ± 28 |
| AGAC | 41.6 ± 1.3 | 56.1 ± 0.3 | 163.2 ± 22.0 | 57.1 ± 0.5 | 1923 ± 52 |
| GAGT | 50.0 ± 2.2 | 64.5 ± 1.2 | 233.9 ± 55.2 | 63.2 ± 2.6 | 2219 ± 34 |
| GAGC | 37.4 ± 0.8 | 62.8 ± 2.3 | 93.5 ± 9.3 | 59.2 ± 2.4 | 2072 ± 24 |
| AAGT | 43.3 ± 1.6 | 57.6 ± 2.0 | 174.5 ± 6.7 | 62.8 ± 1.6 | 1993 ± 15 |
| AAGC | 40.8 ± 0.6 | 69.5 ± 1.4 | 99.1 ± 0.8 | 79.8 ± 3.1 | 2551 ± 52 |
| GAAT | 51.0 ± 2.4 | 57.0 ± 2.0 | 484.7 ± 69.6 | 54.8 ± 4.2 | 1981 ± 25 |
| GAAC | 52.5 ± 2.7 | 58.7 ± 3.1 | 496.5 ± 18.6 | 56.7 ± 2.2 | 2155 ± 97 |
| AAAT | 53.9 ± 0.2 | 59.2 ± 0.9 | 613.0 ± 78.7 | 62.5 ± 1.0 | 2185 ± 8 |
| AAAC | 50.4 ± 0.7 | 59.7 ± 1.6 | 327.7 ± 17.1 | 57.1 ± 2.4 | 2203 ± 47 |
| Average | 41.8 ± 7.4 | 60.1 ± 3.7 | 209.0 ± 170.6 | 61.8 ± 6.4 | 1960 ± 264 |

Table 4: Persistence length $A$, its static $A_s$ and dynamic $A_d$ components and twist $C$ and stretch modulus $B$ of YRRR tetranucleotide sequences.

| Sequence | $A$ $(nm)$ | $A_d$ $(nm)$ | $A_s$ $(nm)$ | $C$ $(nm)$ | $B$ $(pN)$ |
|---|---|---|---|---|---|
| TGGG | 17.1 ± 0.3 | 59.8 ± 1.0 | 24.0 ± 0.5 | 60.6 ± 1.7 | 1673 ± 67 |
| TGGA | 18.0 ± 1.0 | 52.5 ± 0.8 | 27.6 ± 2.4 | 62.0 ± 0.8 | 1695 ± 82 |
| CGGG | 21.5 ± 1.4 | 65.4 ± 0.8 | 32.2 ± 2.9 | 56.6 ± 2.5 | 1974 ± 40 |
| CGGA | 21.9 ± 0.7 | 54.5 ± 0.6 | 36.6 ± 2.0 | 51.6 ± 1.9 | 2000 ± 44 |
| TGAG | 33.2 ± 1.0 | 65.5 ± 2.2 | 67.3 ± 2.2 | 72.7 ± 2.7 | 2335 ± 100 |
| TGAA | 28.0 ± 0.5 | 54.9 ± 1.8 | 57.5 ± 3.1 | 69.0 ± 1.1 | 2044 ± 28 |
| CGAG | 27.6 ± 0.7 | 61.9 ± 0.8 | 50.0 ± 2.3 | 69.3 ± 1.5 | 2377 ± 65 |
| CGAA | 31.6 ± 0.6 | 57.6 ± 2.2 | 70.2 ± 1.0 | 54.3 ± 1.3 | 2357 ± 100 |
| TAGG | 20.1 ± 0.7 | 52.6 ± 1.9 | 32.7 ± 1.2 | 59.8 ± 3.9 | 1663 ± 43 |
| TAGA | 16.8 ± 0.3 | 49.5 ± 0.6 | 25.5 ± 0.8 | 74.1 ± 3.3 | 1593 ± 40 |
| CAGG | 23.0 ± 1.3 | 60.9 ± 1.6 | 36.9 ± 2.9 | 63.5 ± 2.1 | 1996 ± 23 |
| CAGA | 22.4 ± 0.3 | 54.3 ± 0.4 | 38.1 ± 0.8 | 60.5 ± 2.4 | 1958 ± 83 |
| TAAG | 32.2 ± 1.5 | 55.8 ± 1.5 | 76.5 ± 6.9 | 67.5 ± 1.8 | 2186 ± 49 |
| TAAA | 35.0 ± 0.9 | 51.4 ± 2.2 | 110.4 ± 9.5 | 75.7 ± 1.8 | 2161 ± 63 |
| CAAG | 35.1 ± 1.6 | 66.9 ± 2.1 | 73.7 ± 4.9 | 81.2 ± 5.1 | 2566 ± 110 |
| CAAA | 34.6 ± 0.1 | 58.0 ± 0.7 | 85.5 ± 1.3 | 65.1 ± 3.2 | 2060 ± 87 |
| Average | 26.1 ± 6.5 | 57.6 ± 5.2 | 52.8 ± 24.6 | 65.2 ± 7.9 | 2040 ± 276 |

Table 5: Persistence length $A$, its static $A_s$ and dynamic $A_d$ components and twist $C$ and stretch modulus $B$ of RRYR /YRYY tetranucleotide sequences.

| Sequence | $A$ $(nm)$ | $A_d$ $(nm)$ | $A_s$ $(nm)$ | $C$ $(nm)$ | $B$ $(pN)$ |
|---|---|---|---|---|---|
| GGTG | 30.4 ± 0.1 | 57.6 ± 0.8 | 64.1 ± 0.7 | 66.0 ± 2.0 | 2076 ± 68 |
| GGTA | 29.2 ± 0.1 | 52.6 ± 0.6 | 65.8 ± 0.4 | 61.1 ± 0.9 | 1783 ± 53 |
| AGTG | 29.2 ± 0.7 | 59.5 ± 1.3 | 57.5 ± 2.4 | 66.0 ± 3.2 | 1996 ± 74 |
| AGTA | 22.7 ± 2.1 | 51.7 ± 1.5 | 40.6 ± 5.7 | 56.5 ± 3.2 | 1854 ± 13 |
| TGTT | 22.8 ± 0.1 | 58.9 ± 0.1 | 37.2 ± 0.2 | 52.5 ± 1.9 | 1832 ± 28 |
| TGTC | 26.1 ± 1.5 | 53.4 ± 0.7 | 51.3 ± 6.0 | 48.8 ± 1.3 | 2035 ± 114 |
| CGTT | 24.6 ± 0.1 | 57.2 ± 0.1 | 43.1 ± 0.3 | 41.1 ± 0.3 | 1931 ± 10 |
| CGTC | 26.6 ± 2.5 | 57.0 ± 0.6 | 50.7 ± 8.4 | 43.8 ± 1.8 | 2309 ± 108 |
| GGCG | 31.5 ± 0.3 | 65.2 ± 0.3 | 61.1 ± 0.9 | 56.3 ± 0.6 | 2462 ± 27 |
| GGCA | 28.1 ± 0.0 | 63.4 ± 1.3 | 50.4 ± 0.9 | 57.8 ± 0.3 | 2293 ± 41 |
| AGCG | 22.2 ± 3.1 | 56.6 ± 4.6 | 38.8 ± 12.4 | 60.5 ± 3.1 | 2171 ± 149 |
| AGCA | 19.8 ± 1.1 | 63.1 ± 3.6 | 29.0 ± 1.6 | 66.1 ± 6.9 | 2266 ± 46 |
| GATG | 25.8 ± 3.5 | 56.2 ± 0.4 | 49.0 ± 11.4 | 52.4 ± 2.9 | 2142 ± 191 |
| GATA | 20.9 ± 0.5 | 48.5 ± 0.2 | 36.7 ± 1.4 | 65.4 ± 1.3 | 1678 ± 74 |
| AATG | 24.5 ± 0.7 | 56.0 ± 0.2 | 43.6 ± 2.3 | 48.7 ± 1.1 | 1714 ± 43 |
| AATA | 25.8 ± 0.5 | 52.2 ± 1.7 | 50.9 ± 0.3 | 56.2 ± 0.8 | 1799 ± 6 |
| Average | 25.6 ± 3.3 | 56.8 ± 4.4 | 48.1 ± 10.2 | 56.2 ± 7.7 | 2021 ± 229 |

Table 6: Persistence length $A$, its static $A_s$ and dynamic $A_d$ components and twist $C$ and stretch modulus $B$ of RRYY tetranucleotide sequences.

| Sequence | $A$ $(nm)$ | $A_d$ $(nm)$ | $A_s$ $(nm)$ | $C$ $(nm)$ | $B$ $(pN)$ |
|---|---|---|---|---|---|
| GGTT | 38.4 ± 0.4 | 65.8 ± 0.8 | 92.4 ± 0.9 | 54.5 ± 0.1 | 2255 ± 24 |
| GGTC | 44.5 ± 0.1 | 65.6 ± 0.5 | 138.4 ± 3.3 | 65.1 ± 1.2 | 2493 ± 1 |
| AGTT | 48.2 ± 0.7 | 60.8 ± 0.9 | 232.1 ± 3.2 | 56.4 ± 0.4 | 2179 ± 12 |
| AGTC | 50.2 ± 5.8 | 68.3 ± 1.5 | 205.5 ± 65.0 | 62.3 ± 1.9 | 2533 ± 59 |
| GGCT | 42.1 ± 0.3 | 65.2 ± 0.6 | 118.9 ± 3.9 | 55.7 ± 2.3 | 2527 ± 28 |
| GGCC | 44.1 ± 0.4 | 65.3 ± 0.0 | 136.3 ± 4.0 | 60.7 ± 1.6 | 2677 ± 18 |
| AGCT | 41.5 ± 1.8 | 58.8 ± 1.4 | 141.7 ± 12.7 | 55.4 ± 0.5 | 2314 ± 67 |
| GATT | 59.4 ± 0.5 | 70.7 ± 0.3 | 370.7 ± 11.9 | 65.2 ± 2.6 | 2413 ± 49 |
| GATC | 58.1 ± 0.4 | 65.9 ± 0.3 | 495.3 ± 17.6 | 66.7 ± 0.9 | 2339 ± 15 |
| AATT | 61.7 ± 1.0 | 64.9 ± 0.7 | 1267.1 ± 144.2 | 57.9 ± 0.9 | 2209 ± 16 |
| Average | 48.8 ± 7.8 | 65.1 ± 3.2 | 319.8 ± 337.9 | 60.0 ± 4.4 | 2394 ± 154 |

Table 7: Persistence length $A$, its static $A_s$ and dynamic $A_d$ components and twist $C$ and stretch modulus $B$ of YRYR tetranucleotide sequences.

| Sequence | $A$ $(nm)$ | $A_d$ $(nm)$ | $A_s$ $(nm)$ | $C$ $(nm)$ | $B$ $(pN)$ |
|---|---|---|---|---|---|
| TGTG | 17.5 ± 0.9 | 47.4 ± 2.1 | 28.0 ± 3.1 | 54.7 ± 4.5 | 1847 ± 136 |
| TGTA | 17.5 ± 0.4 | 43.3 ± 0.9 | 29.3 ± 0.9 | 64.6 ± 4.3 | 1682 ± 40 |
| CGTG | 19.9 ± 0.1 | 51.3 ± 0.8 | 32.7 ± 0.5 | 54.5 ± 1.5 | 2173 ± 51 |
| CGTA | 19.0 ± 0.5 | 44.0 ± 1.0 | 33.6 ± 2.0 | 50.9 ± 3.5 | 1749 ± 52 |
| TGCG | 18.9 ± 0.3 | 48.8 ± 0.9 | 30.7 ± 0.9 | 50.8 ± 1.8 | 2095 ± 53 |
| TGCA | 15.1 ± 0.1 | 49.7 ± 0.3 | 21.7 ± 0.2 | 69.7 ± 1.1 | 1850 ± 30 |
| CGCG | 25.6 ± 0.7 | 56.4 ± 1.9 | 47.1 ± 2.6 | 52.1 ± 1.0 | 2660 ± 63 |
| TATG | 17.7 ± 0.8 | 43.9 ± 0.3 | 29.8 ± 2.3 | 60.6 ± 0.2 | 1676 ± 12 |
| TATA | 17.6 ± 0.8 | 41.1 ± 0.8 | 31.0 ± 2.8 | 70.1 ± 4.1 | 1672 ± 30 |
| CATG | 16.3 ± 0.5 | 53.2 ± 0.9 | 23.6 ± 0.9 | 60.4 ± 0.2 | 1887 ± 38 |
| Average | 18.5 ± 2.7 | 47.9 ± 4.6 | 30.8 ± 6.5 | 58.9 ± 7.0 | 1929 ± 293 |

Table 8: Persistence length $A$, its static $A_s$ and dynamic $A_d$ components and twist $C$ and stretch modulus $B$ of RYRR/YYRY tetranucleotide sequences.

| Sequence | $A\ (nm)$ | $A_d\ (nm)$ | $A_s\ (nm)$ | $C\ (nm)$ | $B\ (pN)$ |
|---|---|---|---|---|---|
| GTGG | 30.0 ± 0.6 | 56.0 ± 1.7 | 64.9 ± 0.6 | 85.1 ± 1.6 | 2271 ± 131 |
| GTGA | 46.1 ± 0.6 | 61.5 ± 0.2 | 184.6 ± 7.5 | 92.2 ± 1.8 | 2645 ± 14 |
| ATGG | 20.7 ± 1.4 | 59.9 ± 1.2 | 31.7 ± 3.7 | 79.3 ± 0.8 | 1746 ± 203 |
| ATGA | 26.6 ± 1.2 | 59.1 ± 1.1 | 48.7 ± 4.1 | 83.3 ± 0.6 | 1875 ± 54 |
| TTGT | 29.6 ± 1.4 | 61.3 ± 2.1 | 57.3 ± 3.5 | 74.4 ± 5.4 | 1898 ± 31 |
| TTGC | 43.6 ± 2.1 | 67.8 ± 1.3 | 122.6 ± 13.1 | 93.4 ± 1.7 | 2829 ± 164 |
| CTGT | 24.0 ± 0.7 | 59.9 ± 0.4 | 40.1 ± 1.9 | 81.7 ± 2.5 | 1889 ± 52 |
| CTGC | 37.9 ± 0.4 | 62.1 ± 2.4 | 97.7 ± 4.1 | 82.2 ± 6.3 | 2497 ± 87 |
| GTAG | 34.9 ± 2.2 | 54.9 ± 3.0 | 96.1 ± 7.4 | 81.1 ± 4.5 | 2147 ± 148 |
| GTAA | 40.3 ± 2.6 | 54.0 ± 1.3 | 161.7 ± 28.9 | 84.9 ± 1.4 | 2378 ± 111 |
| ATAG | 19.7 ± 0.5 | 55.9 ± 0.5 | 30.4 ± 1.2 | 97.3 ± 2.7 | 1504 ± 23 |
| ATAA | 32.0 ± 3.2 | 54.4 ± 1.0 | 79.7 ± 16.8 | 83.8 ± 3.5 | 1919 ± 58 |
| GCGG | 32.6 ± 0.1 | 65.2 ± 2.1 | 65.5 ± 2.4 | 69.3 ± 6.4 | 2659 ± 116 |
| GCGA | 30.2 ± 0.1 | 62.5 ± 0.3 | 58.5 ± 0.8 | 87.1 ± 1.2 | 2503 ± 66 |
| ACGG | 21.7 ± 0.8 | 60.0 ± 0.9 | 34.0 ± 2.0 | 74.4 ± 1.3 | 2050 ± 116 |
| ACGA | 26.9 ± 0.9 | 62.0 ± 1.5 | 47.4 ± 2.0 | 62.0 ± 1.4 | 2111 ± 48 |
| Average | 31.1 ± 7.7 | 59.8 ± 3.8 | 76.3 ± 44.6 | 82.0 ± 8.7 | 2183 ± 363 |

Table 9: Persistence length $A$, its static $A_s$ and dynamic $A_d$ components and twist $C$ and stretch modulus $B$ of RYRY tetranucleotide sequences.

| Sequence | $A\ (nm)$ | $A_d\ (nm)$ | $A_s\ (nm)$ | $C\ (nm)$ | $B\ (pN)$ |
|---|---|---|---|---|---|
| GTGT | 29.6 ± 1.3 | 55.7 ± 0.7 | 63.2 ± 5.4 | 72.6 ± 1.9 | 1599 ± 47 |
| GTGC | 35.1 ± 0.2 | 53.9 ± 0.8 | 100.8 ± 4.3 | 70.7 ± 1.7 | 2014 ± 38 |
| ATGT | 31.0 ± 0.4 | 56.4 ± 0.6 | 69.0 ± 0.8 | 83.8 ± 3.7 | 1455 ± 54 |
| ATGC | 29.4 ± 1.0 | 59.8 ± 1.7 | 58.5 ± 6.0 | 79.7 ± 3.4 | 1785 ± 106 |
| GTAT | 30.6 ± 0.7 | 50.3 ± 0.0 | 78.4 ± 4.6 | 91.0 ± 0.1 | 1550 ± 22 |
| GTAC | 31.5 ± 0.2 | 49.0 ± 0.1 | 88.0 ± 2.2 | 82.2 ± 3.1 | 1792 ± 78 |
| ATAT | 31.6 ± 0.9 | 51.9 ± 1.1 | 81.0 ± 4.9 | 97.5 ± 3.8 | 1448 ± 22 |
| GCGT | 34.0 ± 0.5 | 56.7 ± 0.7 | 85.0 ± 1.8 | 58.6 ± 1.2 | 2047 ± 37 |
| GCGC | 46.4 ± 1.7 | 57.7 ± 0.8 | 239.7 ± 31.4 | 59.9 ± 1.5 | 2642 ± 90 |
| ACGT | 30.9 ± 1.7 | 54.9 ± 0.7 | 71.2 ± 8.5 | 65.8 ± 1.6 | 1569 ± 41 |
| Average | 33.0 ± 4.8 | 54.6 ± 3.2 | 93.5 ± 50.2 | 76.2 ± 12.3 | 1790 ± 349 |

Table 10: Persistence length $A$, its static $A_s$ and dynamic $A_d$ components and twist $C$ and stretch modulus $B$ of YYRR tetranucleotide sequences.

| Sequence | $A$ $(nm)$ | $A_d$ $(nm)$ | $A_s$ $(nm)$ | $C$ $(nm)$ | $B$ $(pN)$ |
|---|---|---|---|---|---|
| TTGG | 26.7 ± 0.8 | 59.6 ± 2.7 | 48.5 ± 1.1 | 88.0 ± 5.0 | 2313 ± 81 |
| TTGA | 32.1 ± 0.7 | 64.6 ± 2.1 | 64.0 ± 1.5 | 93.7 ± 1.2 | 2466 ± 86 |
| CTGG | 30.8 ± 1.5 | 61.6 ± 1.6 | 61.7 ± 4.6 | 87.3 ± 1.3 | 2529 ± 66 |
| CTGA | 36.0 ± 2.0 | 67.1 ± 2.4 | 78.5 ± 10.5 | 94.4 ± 2.4 | 2597 ± 107 |
| TTAG | 32.9 ± 1.3 | 57.2 ± 1.6 | 77.3 ± 4.2 | 84.4 ± 2.0 | 2245 ± 70 |
| TTAA | 33.7 ± 0.9 | 55.7 ± 0.6 | 85.5 ± 4.3 | 80.4 ± 1.9 | 2276 ± 55 |
| CTAG | 34.5 ± 1.3 | 56.8 ± 1.9 | 87.7 ± 4.1 | 78.7 ± 8.1 | 2365 ± 97 |
| TCGG | 26.3 ± 0.3 | 66.0 ± 1.5 | 43.7 ± 0.9 | 83.3 ± 3.0 | 2546 ± 132 |
| TCGA | 29.6 ± 1.4 | 65.4 ± 0.3 | 54.4 ± 4.5 | 81.1 ± 2.4 | 2698 ± 45 |
| CCGG | 27.4 ± 0.6 | 65.0 ± 0.8 | 47.3 ± 1.5 | 83.0 ± 2.3 | 2755 ± 61 |
| Average | 31.0 ± 3.2 | 61.9 ± 4.1 | 64.9 ± 15.6 | 85.4 ± 5.1 | 2479 ± 168 |

Table 11: Averages and standard deviations of tetranucleotide elastic constants.

| Parameter | Average |
|---|---|
| $A$ $(nm)$ | 31.8 ± 1.0 |
| $A_d$ $(nm)$ | 58.8 ± 5.9 |
| $A'_d$ $(nm)$ | 64.3 ± 7.1 |
| $A_s$ $(nm)$ | 108.1 ± 158.9 |
| $C$ $(nm)$ | 68.0 ± 12.0 |
| $B$ $(pN)$ | 2054 ± 354 |
